# Spatial-DC: a robust deep learning-based method for deconvolution of spatial proteomics

**DOI:** 10.1101/2024.12.18.629082

**Authors:** Yuan Li, Qian Kong, Zihan Wu, Yanfen Xu, Yiheng Mao, Yunjie Gu, Xi Wang, Weina Gao, Ruijun Tian, Jianhua Yao

## Abstract

Spatial proteomics studies the spatial distribution of proteins within cells or tissues, providing a new perspective for comprehending cellular processes and disease mechanisms. One important challenge of current spatial proteomics technologies is low resolution, resulting in multiple cells in each spatial proteomics spot. While methods have been proposed to infer the composition of potential cell types in such spots for spatial transcriptomics, the depressed correlation and divergent quantification between transcriptome and proteome limits their capability in spatial proteomics.

To enhance the utility of spatial proteomics data, we propose Spatial-DC (Spatial Digital Cytometry), a deep learning-based framework that infers cell-type composition in each spot and reconstructs spatially and cell-type resolved proteomic profiles. We achieve this by utilizing transfer learning and self-supervised learning with graph convolutional networks (GCN), which enables the incorporation of target spatial proteomics with reference single-cell or single-cell-type proteomics data.

Through extensive simulations of spatial proteomics data, we demonstrate that Spatial-DC outperforms eight state-of-the-art methods in estimating cell-type composition and provides meaningful reconstructions of proteomic profiles for individual cell types. We apply Spatial-DC to different tissues measured using both multiplexed antibody-based and mass spectrometry (MS)-based spatial proteomics technologies. The results showcase Spatial-DC’s superior sensitivity in providing more refined cell-type distribution maps compared to cell-type-specific marker-based distributions and its feasibility in reconstructing spatially and cell-type resolved proteomic profiles from real-world spatial proteomics data. Moreover, we validate Spatial-DC with a newly self-collected pancreatic cancer spatial proteomics data characterized by complex tumor microenvironment (TME) and identify the signaling direction and strength at both spatial and cell-type levels within the TME, suggesting cell-type-specific and spatially-resolved interactions linked to tumor outcomes.

Our results highlight Spatial-DC as a versatile tool for the deconvolution of spatial proteomics data across various tissue slices, providing a reliable foundation for downstream analysis.

## Introduction

Cell-type resolved composition of tissue microenvironment and cellular communities reflects the physiological state of the tissue^1–4^. For instance, pancreatic cancer composes of various regional sub-tumor microenvironments (subTMEs) characterized by reactive and deserted states, which have discernible impacts on the development and treatment response of tumors^5^. Proteins play a crucial role as direct conductors in the biological processes of life and can serve as effective drug targets^6^. Despite the rise of spatial transcriptomic technologies^7–9^, which have partially resolved some of the aforementioned issues^6,10,11^, the relatively low correlation between transcriptome and proteome limits their further application^12–14^.

Spatial proteomics, an emerging and rapidly advancing technique, has been selected as Method of the Year 2024 due to its pivotal role in profiling the proteome of cells or tissues^15^, thus addressing the challenge of dissecting the organization and functions of complex tissue microenvironment^12,16–18^. Spatial proteomics technologies can be classified into two primary categories: (1) antibody-based spatial proteomics and (2) mass spectrometry (MS)-based spatial proteomics^12,19^. Antibody-based spatial proteomics has excellent throughput of spots, but the depth of detected protein relies on the specificity of antibodies (i.e., spatial-CITE-seq^20^). Spatial MS-based proteomics is an unbiased method for proteome profiling, but it is limited by low spatial resolution (i.e., MASP^21^). A common shortcoming of existing spatial proteomics profiling methods is the lack of single-cell or single cell-type resolution for each spot, highlighting the need for a deconvolution tool to explore the potential of spatial proteomics data.

Over the past decade, various methods have been proposed to deconvolute tissue microenvironments in spatial transcriptomics^22–24^, including: (1) probabilistic model-based methods, such as cell2location^25^, stereoscope^26^, and DestVI^27^; (2) matrix factorization-based methods, such as SPOTlight^28^, CARD^29^ and SpatialDWLS^30^; (3) deep learning-based methods, such as Tangram^31^; (4) canonical correlation analysis-based methods, such as Seurat^32^. However, the significant gap in technology and data distribution between spatial transcriptomics and spatial proteomics limits the potential for generalizing these methods to spatial proteomics, especially when many of them were designed based on certain specific distributions of spatial transcriptomics. For example, cell2location^25^ is a Bayesian model that decomposes the spatial expression matrix into a predefined set of cell type signatures derived from a reference single-cell or single-nucleus expression profile. In this process, the input matrices are modeled as following negative binomial distribution. The same assumption of data distribution is applied in stereoscope^26^ and DestVI^27^. These assumptions are strongly correlated with spatial transcriptomics but do not align with the characteristics of spatial proteomics data.

In addition, some algorithms proposed for the deconvolution of spatial transcriptomics rely on selected informative genes from reference data and spatial transcriptomics data, usually consisting of highly variable genes or cell-type-specific genes. For example, Seurat^32^ uses a ranked list of highly variable genes to identify shared anchors between reference data and spatial transcriptomics data, and then assigns cell type labels to the target data based on these shared anchors. SPOTlight^28^ conducts deconvolution using non-negative matrix factorization regression (NMF) and non-negative least squares (NNLS), which relies on intersected cell-type-specific genes from both reference data and spatial transcriptomics data. CARD^29^ builds upon a non-negative matrix factorization model using informative genes from reference data and accounts for the spatial correlation structure by utilizing a conditional autoregressive (CAR) model. SpatialDWLS^30^ is developed for deconvolution based on dampened weighted least squares (DWLS), which depends on a set of signature genes from reference data. However, the performance of these deconvolution approaches may be affected by technical and biological biases introduced by the selected informative genes^33,34^. It could be beneficial for both spatial transcriptomics and spatial proteomics to partially eliminate these biases by designing methods based on entire expression profiles^34,35^.

Moreover, the presence of multiple cell types within each spot results in a detected proteomic expression profile comprising a mixture of the expression profiles from these individual cell types. The absence of proteomic expression profile information for each individual cell type limits further analysis on a specific cell type or interactions between certain cell types. Despite obtaining the composition of cell types in each spot, the proteomic expression profiles of each individual cell type cannot be directly reconstructed. Hence, in addition to cell-type composition, the reconstructions of the proteomic profiles for each cell type within a spot is also crucial. Only a very few methods designed for spatial transcriptomics, such as Cottrazm^36^, and some methods for bulk-level non-spatial transcriptomics, such as CIBERSORTx^37^ and TAPE^35^, estimate both the cell-type composition and the reconstructions of cell-type proteomic profiles simultaneously. However, these methods are ineffective or even unusable for spatial proteomics.

To address the aforementioned problems, we present Spatial-DC (Spatial Digital Cytometry), a deep learning-based method designed to predict the cell-type composition and reconstruct the proteomic profiles of cells from spatial proteomics data obtained from various tissues under different kinds of spatial proteomics techniques. Firstly, Spatial-DC trains a distribution model on reference single-cell or single-cell-type proteomics data (reference data) to learn the implicit correspondence between proteomic profiles and cell-type resolved information. Then, the trained distribution model is applied to spatial proteomics data to generate intermediate prediction, including cell-type composition and cell-type proteomic profile weights of each spot. Next, a self-supervised model of graph convolutional networks (GCN) takes the intermediate prediction, along with the spatial proteomic profiles and spatial coordinates of spatial proteomics data, to produce refined prediction of cell-type composition and cell-type proteomic profile weights. Finally, Spatial-DC integrates the spatial proteomic profiles with the refined cell-type proteomic profile weights to obtain the proteomic profiles of individual cell types for each spot.

The performance of Spatial-DC in predicting cell-type composition was evaluated using multiple datasets, including synthetic data and publicly available data from different spatial proteomics techniques under various tissues (normal and disease). The results demonstrated that Spatial-DC outperformed other state-of-the-art methods. We also analyzed the authenticity of the reconstructed spatially and cell-type resolved proteomic profiles generated by Spatial-DC from qualitative, quantitative, and functional enrichment perspectives. In addition, we applied Spatial-DC to a newly self-collected pancreatic cancer spatial proteomics data and validated its predictions using multiplexed immunohistochemical (mIHC). We found a high degree of consistency between the predicted and actual results.

Furthermore, we analyzed cell-cell communications within the TME of pancreatic cancer using the reconstructed spatially and cell-type resolved proteomic profiles produced by Spatial-DC. Our results suggest a potential interaction between *ITGA2* and *COL8A1* at the spatial level, with significant expression observed in pancreatic cancer cells (PC) and cancer-associated fibroblasts (CAF), respectively. Finally, we investigated the association between the co-expression levels of the protein pair *ITGA2* and *COL8A1* and patient outcomes utilizing proteomics data and corresponding clinical information from LinkedOmicsKB^38^. Our analysis demonstrated a strong correlation between the co-expression levels of *ITGA2* and *COL8A1* and the survival time of pancreatic cancer patients, highlighting the potential clinical significance of our findings.

## Results

### Spatial-DC: a deep learning-based method for deconvolution of spatial proteomics

Spatial-DC is a deep learning-based deconvolution method for spatial proteomics data, which transfers cell-type resolved information from reference data to target spatial proteomics data, enabling the inference of cell-type composition as well as the reconstruction of cell-type resolved proteomic profiles for individual spatial spots within a given tissue (**Fig. 1a** left). Spatial-DC offers a comprehensive set of features that empower researchers to: (1) estimate the distribution of different cell types with spatial context; (2) perform spatial domain recognition, identifying distinct regions or patterns in the tissue; (3) discover cell-type-specific proteins that exhibit spatial patterns, providing insights into their functional roles and interactions; and (4) analyze spatially informed cell-cell communications, unraveling the complex networks of interactions within the tissue microenvironment (**Fig. 1a** right).

**Fig. 1.**
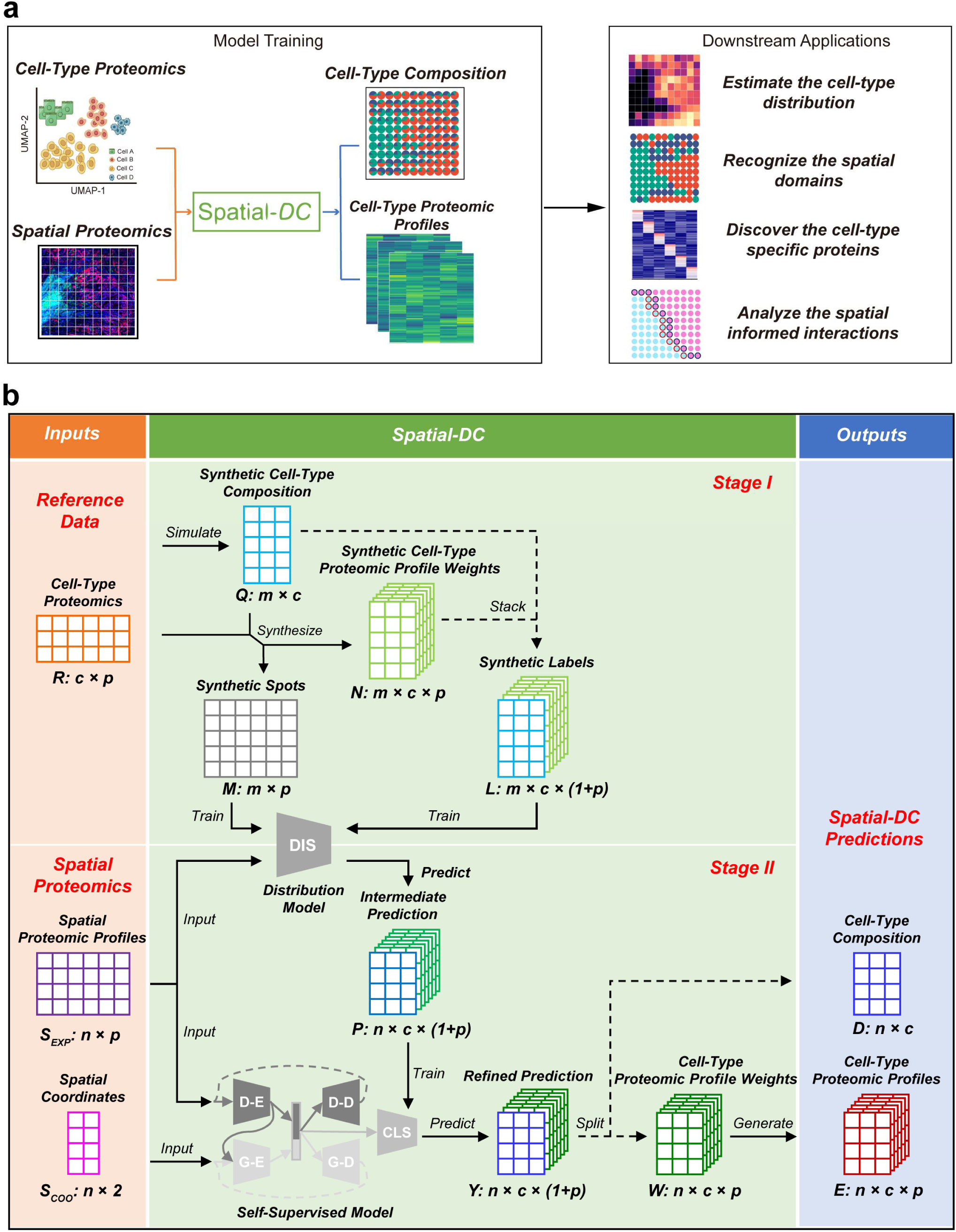
Workflow of Spatial-DC. **a,** A schematic graph for Spatial-DC (left) and subsequently downstream applications (right). **b,** The framework of Spatial-DC, corresponding to the left side of sub-figure **a**. In stage I, Spatial-DC requires reference proteomics data ***R*** (e.g., cell-type proteomics) as input and obtains synthetic cell-type composition ***Q*** from a Dirichlet distribution (**Methods**). ***R*** and ***Q*** are then used to generate synthetic spots ***M*** and synthetic cell-type proteomic profile weights ***N***. Subsequently, ***Q*** and ***N*** are stacked to produce synthetic labels ***L***. ***M*** and ***L*** are used as training data and labels, respectively, to train a deep neural network-based distribution model **DIS**. In stage II, Spatial-DC utilizes spatial proteomics data, including spatial proteomic profiles ***S_EXP_*** and associated spatial coordinates ***S_COO_*** as inputs. ***S_EXP_*** are fed into the trained **DIS** model to produce intermediate prediction ***P***. Subsequently, ***S_EXP_***, ***S_COO_*** and ***P*** are utilized to train a self-supervised model **SSM**, which is composed of a variation graph autoencoder (**VGAE**), a deep autoencoder (**DAE**) and a classifier (**CLS**) (**Fig. S1**). Once trained, the model produces refined prediction ***Y*** from the classifier **CLS**, which are then split into cell-type composition ***D*** and cell-type proteomic profile weights ***W***. By using ***W***, along with the spatial proteomic profiles ***S_EXP_***, cell-type proteomic profiles ***E*** of each spot are obtained. The abbreviations of the letters in this framework: ***c*** is the number of cell types from reference data ***R***; ***p*** is the number of proteins shared between reference data ***R*** and spatial proteomic profiles ***S_EXP_***; ***m*** is the number of synthetic spots; ***n*** is the number of spots from spatial proteomics ***S_EXP_***; **D-E/D-D** represents the encoder and decoder architecture of autoencoder; **G-E*/*G-D** represents the encoder and decoder architecture of VGAE.

As a deep-learning based framework for deconvolution of spatial proteomics data, Spatial-DC comprises two stages: stage I involves generating synthetic spots data from reference data to train a cell-type distribution model. In stage II, the trained distribution model is used to produce intermediate prediction of cell-type resolved information from spatial proteomics data, which was subsequently refined based on the characteristic of spatial proteomics data with a self-supervised model and ultimately transformed into cell-type composition and cell-type proteomic profiles (**Fig. 1b**). Technically, the distribution model in stage I is a deep neural network (DNN) composed of three stacked fully connected layers, which is trained with the reference data to learn the implicit relationships between proteomic profiles and cell-type resolved information. In stage II, the intermediate prediction of cell-type resolved information of each spot is initially estimated by passing the spatial proteomics data through the trained distribution model from stage I. Then, a self-supervised model comprising a variation graph autoencoder (VGAE), a deep autoencoder and a classifier is trained to integrate the intermediate prediction of cell-type resolved information with the expression and spatial characteristic of the spatial proteomics data. The VGAE and the deep autoencoder produce embeddings of the expression profiles and spatial information of the spatial proteomics data, respectively. The classifier then establishes the connection between these embeddings and the cell-type resolved information. The two autoencoders are trained using a self-supervised strategy with two reconstruction losses, while the classifier is trained with the intermediate cell-type resolved information through a supervised learning procedure. The well-trained self-supervised model is finally re-applied on the target spatial proteomics data to obtain the cell-type composition as well as cell-type resolved proteomic profiles of each spot (**Fig. 1b; Methods; Fig. S1 and Table S1**).

### Assessment of Spatial-DC for estimating cell-type composition and reconstructing cell-type proteomic profiles on synthetic spatial proteomics data

The assessment of spatial deconvolution methods frequently necessitates the utilization of synthetic spatially-resolved data^22–24^. These synthetic datasets incorporate a predefined ground truth and preserve the relative spatial patterns and structures from annotated spatially and single-cell resolved tissue slices, allowing for an unbiased and objective evaluation of the performance of a given deconvolution method^39–41^.

To systematically assess the deconvolution performance of Spatial-DC, we developed a framework to generate synthetic spatial proteomics data (**Fig. 2a; Methods; Fig. S2a**). Briefly, we created a comprehensive synthetic dataset of spatially and single-cell resolved proteomics data for human non-small-cell lung cancer (NSCLC) (**Fig. 2b; Table S2**), which was subsequently utilized to generate synthetic spatial proteomics data encompassing various conditions, for the comprehensive assessment of Spatial-DC. We first acquired two distinct MS-based single-cell proteomics datasets from separate sources^42,43^, which have 1,003 shared proteins and both encompass three cell types: macrophage cell line (RAW), a respiratory epithelial cell line (C10) and an endothelial cell line (SVEC). One dataset^42^, collected from nanowell chip based single-cell proteomics, was utilized to simulate spatial proteomics data and is referred to as NSCLC projection data. Another dataset^43^ collected from nanoPOTS chip based single-cell proteomics, served as the reference for deconvolution and is referred to as NSCLC reference data (**Fig. S2b,S2c**). Noted that we used a separate proteomics dataset as a reference to deconvolute the synthetic spatial proteomics data. This is crucial, as it necessitates accounting for real-world difference (such as cross-subject and batch-related variances) between the reference and projection data^24^. Besides, we leveraged the precisely annotated, single-cell resolved NSCLC spatial structure data from the CoxMx SMI platform (www.nanostring.com) to mimic the complex distribution of tissue niches (**Fig. S2d**). The synthetic NSCLC spatially and single-cell resolved proteomics data was derived from the integration of spatial structure data and projection data. Specifically, we used the NSCLC spatial structure as a reference for cell locations and replaced each cell with its corresponding proteomic profiles from the NSCLC projection data (**Methods**). Using this synthetic NSCLC spatially and single-cell resolved proteomics data, we were able to simulate spatial proteomics data with varying spot sizes along with corresponding cell-type composition for each spot. And to simulate complex and diverse states as in real-world datasets, external Gaussian noise was added to the synthetic spatial proteomics data (**Methods**).

**Fig. 2.**
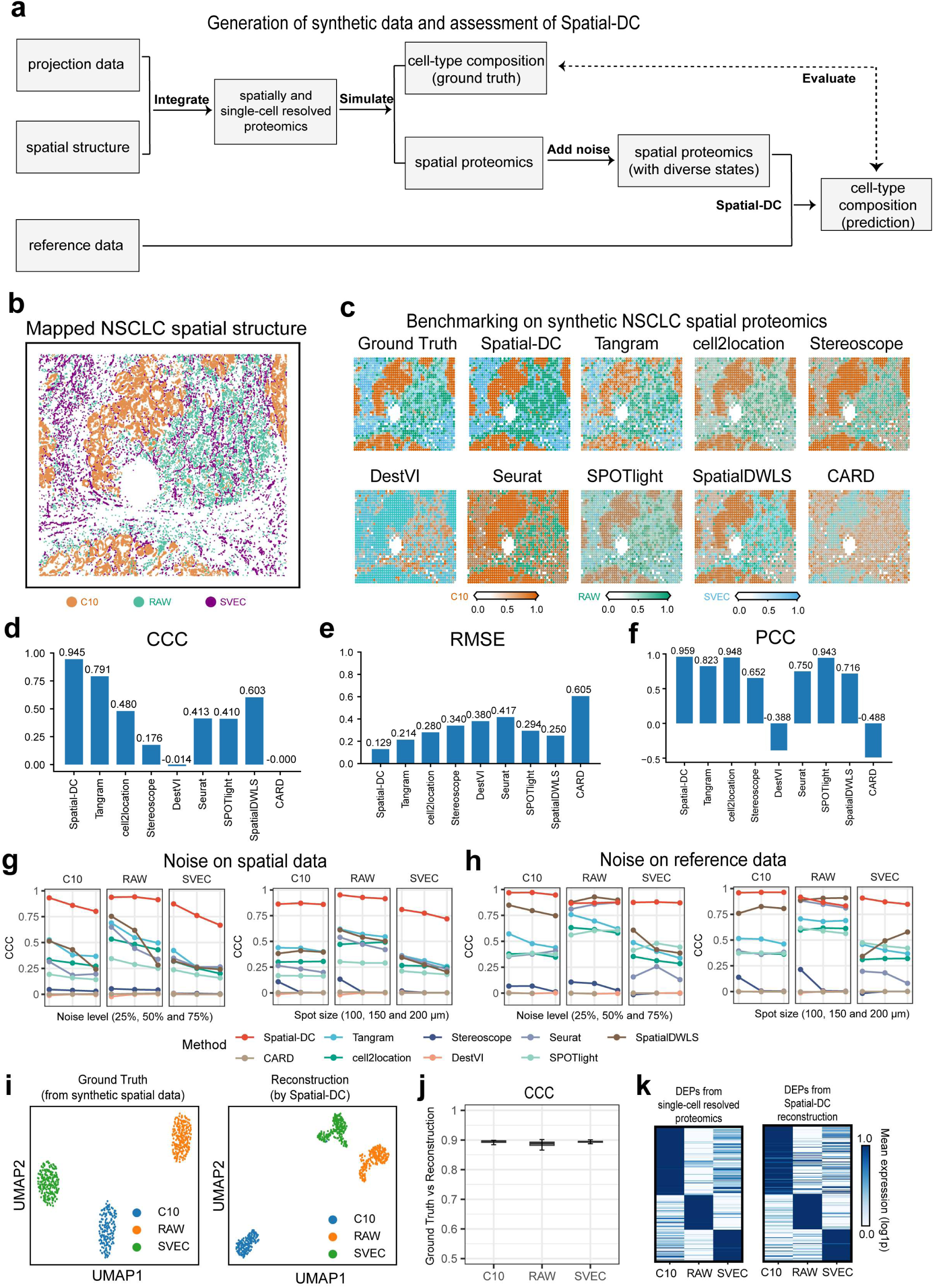
Assessment of Spatial-DC on synthetic NSCLC data. **a,** The typical framework of generating synthetic spatial proteomics data, defining a well-established cell-type composition, and evaluating Spatial-DC with another single-cell resolved proteomics data as a reference (**Methods**). **b,** Spatial map illustrating the distribution of cells resolved at a single-cell level, derived from synthetic NSCLC spatially and single-cell resolved proteomics data, encompassing 17,837 C10 cells, 7,616 SVEC cells, and 7,409 RAW cells. **c-f,** Performance evaluation of Spatial-DC and eight state-of-the-art deconvolution methods, including Tangram^31^, cell2location^25^, Stereoscope^26^, DestVI^27^, Seurat^t32^, SPOTlight^28^, SpatialDWLS^30^ and CARD^29^ on representative synthetic NSCLC data (with spot size of 100 μm), where each spot contains an average of 17 cells without external Gaussian noise added (**Methods; Table S4**). **c,** Representative spatial maps depicting the distribution of cell types on ground truth and predictions. **d-f**, Quantitative assessment of Spatial-DC on representative synthetic NSCLC data, which encompasses CCC, PCC, and SPCC metrics, by comparing cell-type composition between the ground truth and the predictions (**Methods**). Each bar represents the mean value of the metrics from three cell types for a given method. **g-h,** Evaluation of the robustness of Spatial-DC on synthetic NSCLC data. Line plots showing the CCC evaluation metrics of Spatial-DC and the eight state-of-the-art deconvolution methods across three cell types (left, middle and right) under varying noise levels (left column) and different spot sizes (right column), subject to the noise perturbed spatial data (**g**) and reference data (**h**), respectively (**Methods**). Each data point, corresponding to varying noise levels (left column), represents the mean CCC metrics from three spot sizes, and vice versa. **i-k**, Assessment of reconstructing cell-type proteomic profiles by Spatial-DC from representative synthetic NSCLC data (with spot size of 200 μm). **i,** UMAP visualizations depicting 647 proteomic profiles from the single-cell-type resolved samples, encompassing a total of 1,003 proteins for both. These proteomic profiles comprise three NSCLC cell types—183 C10 cells, 241 RAW cells, and 223 SVEC cells—derived from the ground truth (**Fig. S2a**) and the reconstructions by Spatial-DC. **j,** Quantitative assessment of reconstructed cell-type proteomic profiles by Spatial-DC from the synthetic NSCLC data. **k,** Heat maps visualizing the log1p-normalized proteins abundance for NSCLC projection data^43^ and NSCLC reconstructed data by Spatial-DC using differentially expression proteins (DEPs) panel, consisting of 103 proteins for C10 cells, 52 proteins for RAW cells, and 48 proteins for SVEC cells. These protein panels were identified in both NSCLC projection^42^ and reference data^43^ (**Methods**).

Using synthetic NSCLC spatial proteomics data, we evaluated Spatial-DC and eight state-of-the-art deconvolution methods, including Tangram^31^, cell2location^25^, Stereoscope^26^, DestVI^27^, Seurat^t32^, SPOTlight^28^, SpatialDWLS^30^ and CARD^29^ (**Methods and Table S4**). Our benchmarking analysis on cell-type composition prediction revealed that Spatial-DC yielded results more closely aligned with the ground truth compared to the other eight methods. Specifically, Tangram, cell2location, Seurat, SPOTlight and SpatialDWLS demonstrated inferior performance, while Stereoscope, DestVI and CARD were unable to process the synthetic spatial proteomics data in our experiments (**Fig. 2c**). The advantage of Spatial-DC was further substantiated by various evaluation metrics, including Concordance Correlation Coefficient (CCC), Root Mean Squared Error (RMSE), Pearson Correlation Coefficient (PCC) and Spearman Correlation Coefficient (SPCC) (**Fig. 2d-f; Fig. S2e and Table S4; Methods**). Then, we thoroughly evaluated the robustness of Spatial-DC by intentionally introducing varying degrees of Gaussian noise (25%, 50% and 75%) to the synthetic spatial proteomics data simulated with different spot sizes (100, 150 and 200 μm). The same levels of noise were also introduced into the reference data for evaluation. Moreover, external evaluation studies were conducted under different perturbed conditions for both reference and spatial proteomics, involving joint Gaussian noise and number of highly variable proteins retained for deconvolution (**Methods**). Our results demonstrated that Spatial-DC consistently exhibited the most resilient prediction capability and outperformed the other deconvolution across difference conditions, highlighting its robustness for a wide range of spatial proteomics applications (**Fig. 2g,h; Fig. S3-S5 and Table S4**).

Then, we sought to evaluate the performance of reconstructing cell-type proteomic profiles of Spatial-DC on synthetic NSCLC data (**Methods**). Employing UMAP analysis, we found that the three distinct cell types—C10, RAW and SVEC cells—were clearly segregated in both the ground truth data and the reconstructed data generated by Spatial-DC (**Fig. 2i**). This observation underscored the ability of Spatial-DC to preserve the underlying cell-type identities. To provide a quantitative assessment of the reconstructed cell-type proteomic profiles, we calculated CCC metrics, together with PCC, SPCC and RMSE metrics, between the scaled proteomic profiles of each spot from the synthetic NSCLC spatial proteomics data and those reconstructed by Spatial-DC. The results revealed a high degree of correlation, indicating the accurate reconstruction of Spatial-DC (**Fig. 2j; Fig. S6**). Additionally, we performed statistical analyses on both projection data and reference data to ascertain significance of individual proteins within each cell type (**Methods**). Following this, we selected and retained those proteins that were significantly expressed as cell-type-specific markers. Comparing the reconstruction of Spatial-DC with the ground truth using these cell-type-specific markers, we observed a remarkable degree of similarity, thereby validating the efficacy of our approach in discovering potential cell-type-specific markers after reconstruction (**Fig. 2k and Table S4**).

Finally, we conducted several ablation studies on Spatial-DC using synthetic NSCLC data, which investigated various components or parameters, including the inclusion of Stage II, the number of simulated spots utilized for training distribution model, and the number of nodes used to construct the nearest neighbor graph. The results demonstrated that Spatial-DC is robust and consistently delivers high performance across a wide range of parameter settings (**Fig. S7 and Table S4**).

In summary, our experiments on synthetic spatial proteomics data have shown that Spatial-DC excels in deconvolution, especially when confronted with varying levels of external noise and heterogeneous cell distributions across different tissue states. Moreover, the extensive evaluation on synthetic data underscored Spatial-DC’s proficiency in precisely reconstructing proteomic profiles for distinct cell types, thereby enhancing its potential to provide profound insights into the intricate spatial organization and functional roles of cells within complex biological tissue environments.

### Application of Spatial-DC on human palatine tonsil data from spatial antibody-based proteomics technology

To assess the deconvolution capability of Spatial-DC on real-world spatial proteomics data encompassing diverse detection technologies and tissue states, we curated a comprehensive dataset and conducted in-depth experiments (**Table S3 and Note S1**). Initially, we focused on human palatine tonsil data sequenced using antibody-derived DNA tags (ADTs)^20,44^. The palatine tonsil, a secondary lymphoid tissue, harbors an intricate immunological microenvironment characterized by follicles separated by an inter-follicular zone^45^. Germinal centers (GCs) reactions initiate within the center of these follicles, pivotal for humoral immunity against pathogens^20,44^. The reference proteomics data encompasses a total of 17,367 tonsillar cells, categorized into three primary groups: B cells, T cells and stromal cells (**Fig. 3a**). The B cell population is further subdivided into five distinct subtypes: naïve B cells (NBC), activated naïve B cells (ANBC), germinal center B cells (GCBC), memory B cells (MBC) and plasma cells (PC). The T cells consist of six subtypes, including naïve CD4 T cells (Naïve T4), T4, naïve CD8 T cells (Naïve T8), T8, cycling T cells (Cycling T) and double-negative T cells (DN). The stroma cell category encompasses eight subtypes: dendritic cells (DC), plasmacytoid DC (PDC), follicular DCs (FDC), monocytes (MO), granulocyte cells (GN), epithelial cells (Epi), natural killer cells (NK), and innate lymphoid cells (ILC). These cellular populations collectively contribute to and maintain the majority of functional events that occur within the human palatine tonsil^44,46^. The spatial proteomics data consists of 282 proteins identified from a 2.5 mm × 2.5 mm slice of a palatine tonsil, with spot sizes of 25 μm. Through the analysis of processed bright-field image of the tonsil slice, we discerned distinct spatial structures within the tissue (**Fig. 3b**). The palatine tonsils consist of three representative compartments: the crypt zone, the inter-follicular zone, and the follicles^45,47^, which are enriched with epitheliums, T cells, and B cells, respectively. We applied unsupervised clustering with the k-means algorithm to the palatine tonsil data and revealed the three distinct spatial domains (**Fig. 3c; Methods**), which also align with the enriched expression of known cell-type-specific markers^45^, specifically *CDH1* for epitheliums, *CD3* for T cells and *CD19* for B cells (**Fig. S8a**). With the assistant of the reference single-cell proteomics data^44^, we then employed Spatial-DC for the deconvolution of the human palatine tonsil spatial proteomics data. Our deconvolution results revealed that B cells are predominantly found in follicles, while T cells predominantly reside in the inter-follicular zone. Additionally, the crypt zone is primarily populated by Epi and MO (**Fig. 3d; Fig. S8b and Table S5**). These conclusions align with previously reported findings^45,48^.

**Fig. 3.**
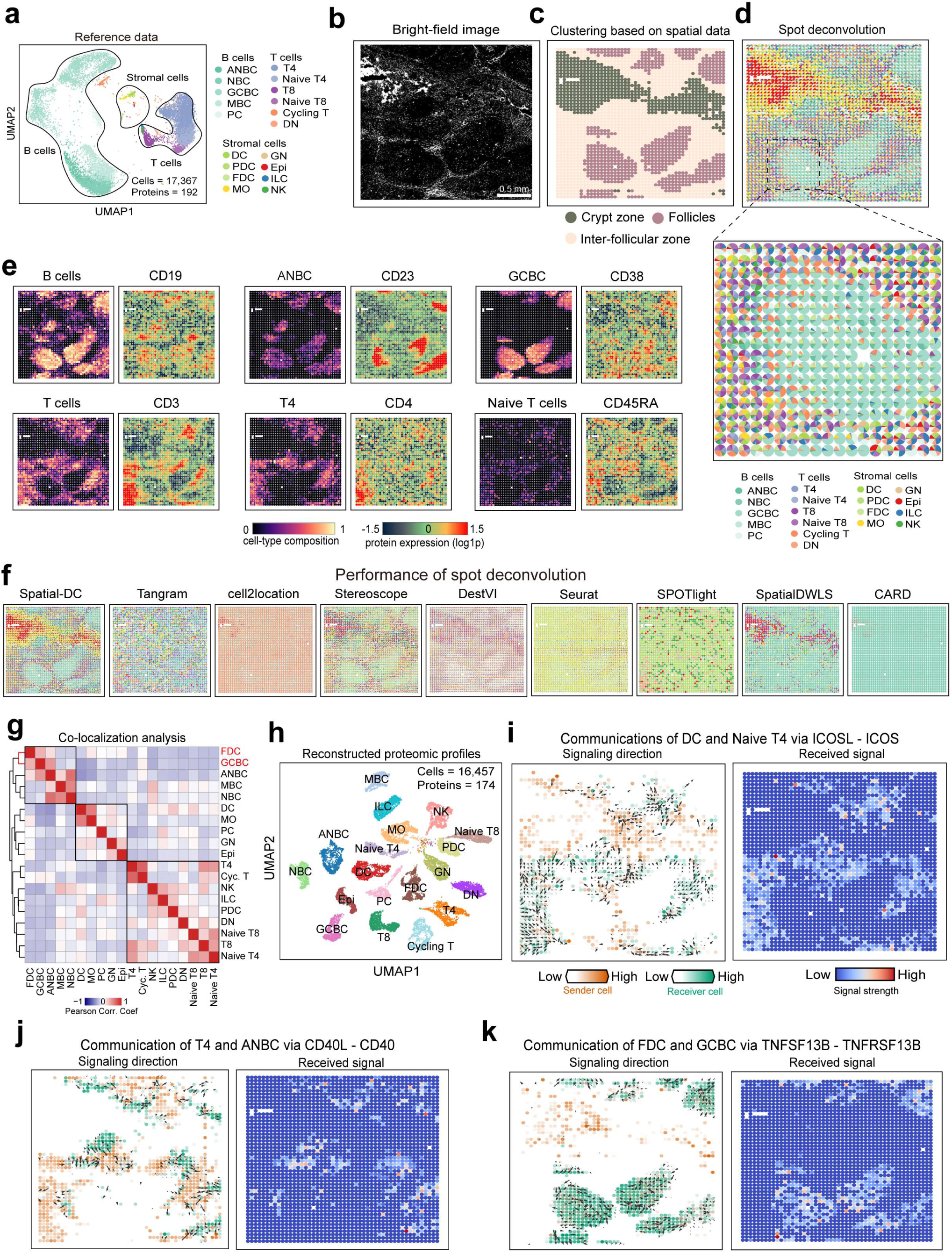
Application of Spatial-DC on human palatine tonsil data from antibody-based proteomics technology. **a-d,** Overview of the human palatine tonsil data. **a,** UMAP visualization of the 17,367 tonsillar cells across 19 cell types sourced from reference data^44^. **b,** Processed bright-field image^20^ of a tonsil slice, related to target tonsil spatial proteomics data (**Methods**). **c,** Spatial map of three domains recognized using spatial data via k-means clustering (k = 3) (**Methods**). **d,** Spatial pie plot of the tonsillar cell-type composition deconvoluted by Spatial-DC. Selected regions have been zoomed in to clearly display the composition in the form of pie charts. **e,** Visualization of cell-type distributions predicted by Spatial-DC (left) and corresponding cell-type-specific markers identified from tonsil spatial proteomics data (right) for six representative subtypes. **f,** Assessment of deconvolution performance of Spatial-DC and eight competing methods, represented as spatial pie plots. **g,** Heat map showing the hierarchical distance of Pearson correlation coefficient calculated from the cell-type composition deconvoluted by Spatial-DC (**Methods**). Three primary patterns exhibiting close distances have been outlined with black squares. **h**, UMAP plot showing the distribution of reconstructed cell-type proteomic profiles by Spatial-DC. **i-k**, The predicted signaling directions and corresponding strength of human palatine tonsil tissue (**Methods**). The signaling directions are represented by black arrowheads, with the cell-type composition of sender cells and receiver cells as the background (left). The strength of the received signals of each spot are shown on the right (right).

Subsequently, we conducted benchmarking studies on the palatine tonsil data to evaluate the performance of Spatial-DC in cell-type composition prediction, comparing it with eight other competing methods (**Methods**). Our results demonstrated that the predicted cell-type composition by Spatial-DC exhibited the closest alignment with established cell-type specific markers (**Fig. 3e; Fig. S8b-S8d; Table S5 and S10**). Besides, Spatial-DC and Stereoscope were the only methods that successfully identified B cells enriched in follicles, whereas the remaining methods failed to make such predictions (**Fig. 3f; Table S5**). These observations further underscored Spatial-DC in accurately deciphering the cellular landscape within the tissue, thereby highlighting its superiority among the evaluated methods.

The establishment of humoral immunity in the tonsils relies on the GCs reaction, which composed of diverse cellular distribution patterns and intricate cellular interactions during the initiation and subsequent maturation of the GCs^49^. To gain further insight of GCs reaction from palatine tonsil, we firstly performed the identification of co-localization patterns using predicted cell-type composition by Spatial-DC (**Methods**) and found three primary patterns (**Fig. 3g**). Notably, within the co-localization pattern comprising five cell types—FDC, GCBC, ANBC, MBC and NBC—FDC exhibited a closer proximity to GCBC than to the other three B cells (**Fig. 3g**). This finding is consistent with their intricate interactions during the formation and maintenance of GCs^50^. Next, we analyzed the reconstructed spatially and cell-type resolved proteomic profiles from the spatial palatine tonsil data by Spatial-DC (**Fig. 3h**). Our analysis identified several hallmark cell-type-specific marker proteins. For instance, we observed *CD19* as a characteristic marker for pan-B cells, *CD3* for pan-T cells, *CD4* specifically in T4 cells, and *CD8* in T8 cells (**Fig. S8e**). Then, we inferred cell-cell communications through known protein-protein interactions (PPIs) from the reconstructed spatial proteomics data using COMMOT^51^ (**Table S5 and Methods**). During the initiation phase of GCs reaction, T cells and B cells undergo activation upon encountering their cognate antigen. Subsequently, activated T cells and B cells engage in prolonged interactions within the inter-follicular zone^49^. Our analysis of the identified PPIs from the reconstructed data corroborated the known interactions between DC and naïve T4 through *ICOSL*–*ICOS*, as well as between T4 and ANBC through *CD40L*–*CD40* within this phase (**Fig. 3i,j**)^49^. As activated T cells and B cells migrate towards the center of the follicles, the early GCs form, characterized by a dense network of FDC^49^. Our results indicated that within these follicles, FDC interact with GCBC through the *TNFSF13B*–*TNFRSF13B* protein pair (**Fig. 3k**). These analyses confirmed the reliability and effectiveness of cell-type-specific proteomic profiles reconstructed by Spatial-DC and highlighted its potential application in inferring the underlying cellular interactions based on these reconstructions.

The present findings not only affirmed the exceptional performance of Spatial-DC beyond eight other deconvolution methods in accurately deconvoluting ADTs-derived spatial proteomics data, but also demonstrated its proficiency in uncovering the precise location of cell types and inferring intricate cellular interactions using reconstructing proteomic signatures, underscoring the broad applicability and potential of Spatial-DC in the analysis of real-world spatial proteomics datasets.

### Application of Spatial-DC on mouse brain coronal data from spatial MS-based proteomics technology

Next, we acquired spatial proteomics data from a coronal section of the mouse brain utilizing micro-scaffold followed by LC-MS/MS^21^, as well as a reference proteomics dataset of mouse cell types obtained via magnetic-activated cell sorting (MACS) followed by LC-MS/MS^52^ (**Table S3; Note S1**). The mouse brain coronal dataset comprises 208 sample slices acquired at low spatial resolution (400 × 400 μm^2^) and includes a total of 5,019 identified proteins, categorized into four distinct spatial domains: cortex (CTX), hippocampus (HIP), thalamus (TH) and hypothalamus (HY) (**Fig. 4a**). The reference proteomics data encompasses neurons and glial cells, the latter of which includes three cell types: astrocytes, oligodendrocytes, and microglia, with each cell type represented by three biological replicates (**Fig. 4b**). Our deconvolution results demonstrated that the cell-type composition generated by Spatial-DC showed high correlations with the representative known cell-type-specific markers (**Fig. 4c-e; Table S6 and S10**).

**Fig. 4.**
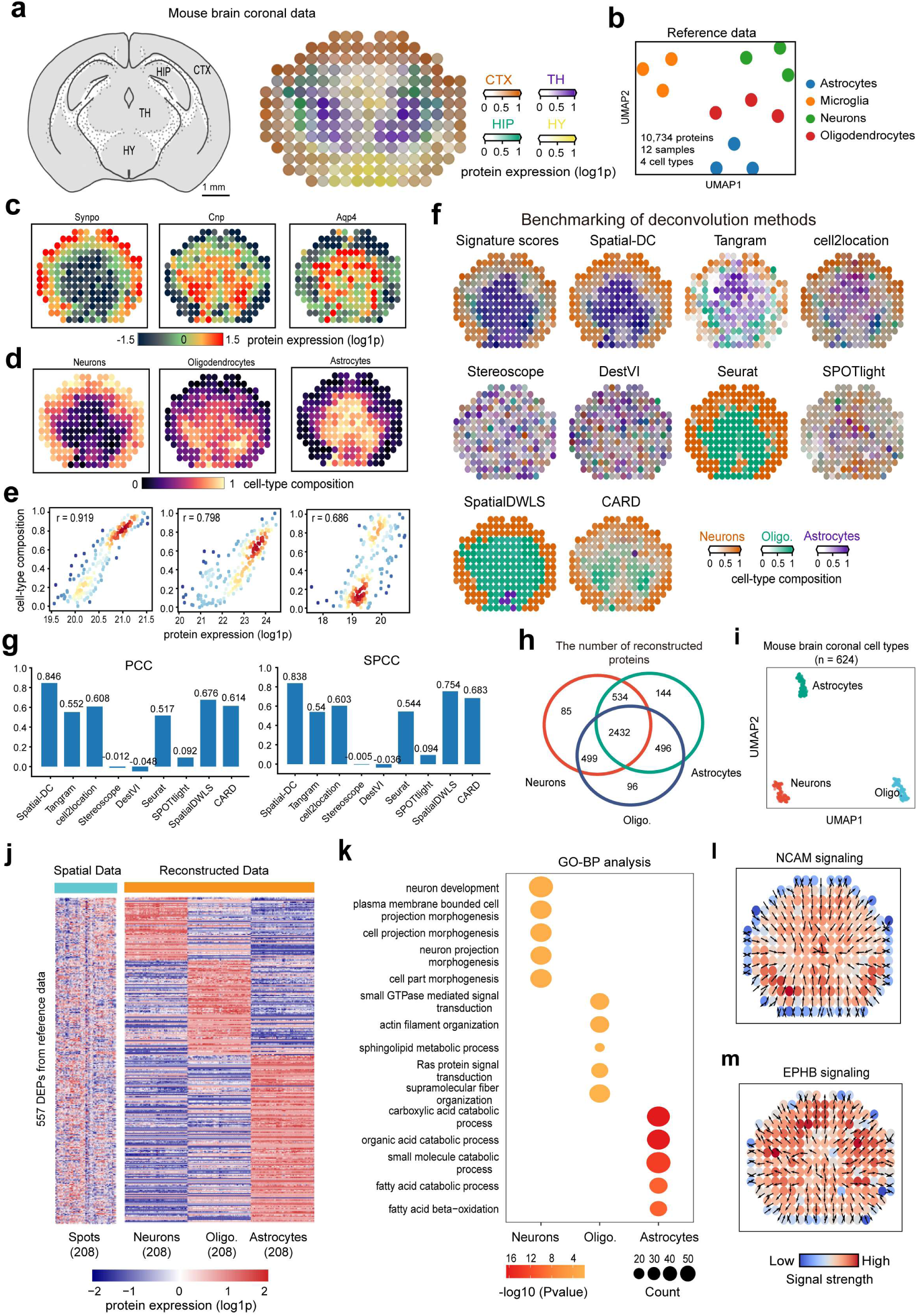
Application of Spatial-DC on mouse brain coronal data from MS-based proteomics technology. **a-b,** Overview of the mouse brain coronal data. **a,** The tissue structures of a mouse brain coronal section^70^, highlighting four primary anatomical regions (left), and the annotated distribution of these regions using established protein panels (right). **b,** UMAP visualization of the reference MS-based mouse brain data^52^. **c-e,** Quantitative assessment of Spatial-DC’s deconvolution performance on mouse brain coronal data. **c,** Spatial maps of representative known cell-type-specific markers from mouse brain coronal data (*Synpo* for neurons, *Cnp* for oligodendrocytes, and *Qqp4* for astrocytes). **d,** The predicted cell-type composition (ranging from 0 to 1) of mouse brain coronal data by Spatial-DC. **e,** Scatter plots comparing the linear correlation between identified representative cell-type-specific marker protein abundance in **c** and predicted cell-type composition in **d** for three cell types, respectively. **f-g,** Benchmarking study comparing signature scores with cell-type composition predicted by Spatial-DC and eight competing methods. Signature scores were obtained using a panel of highly correlated cell-type-specific markers (**Methods**). The visualizations are shown in **f** and the corresponding metrics (PCC and SPCC) are represented as bar plots in **g**. **h-i**, Evaluation of reconstructed cell-type proteomic profiles by Spatial-DC. **h**, Venn plot showing the reconstructed proteins for three cell types. **i,** UMAP plot of the reconstructed data by Spatial-DC. **j,** Comparison of 557 differentially expressed proteins (DEPs) from original spatial proteomics data (left) and reconstruction by Spatial-DC (right). The DEPs were calculated using one-way ANOVA analysis based on mouse brain reference data. **k,** Dot plot exhibiting the 15 significantly enriched GO-BP terms for the three cell types using cell-type-specific proteins. **l-m**, The predicted signaling directions and corresponding strength of the NCAM and EPHB pathways from the reconstructed data of mouse brain coronal (**Methods**).

To evaluate the deconvolution performance of Spatial-DC and eight comparative methods in the absence of definitive ground truth for the primary cell types of the mouse brain coronal data, we devised a surrogate approach utilizing the abundance of highly correlated and cell-type-specific markers, referred to as signature scores (**Fig. 4f; Methods; Fig. S9a and Table S6,S10**). Our comparative evaluation analysis between signatures scores and deconvolution outcomes indicated that Spatial-DC predicted the cell-type distribution with the highest degree of similarity to those defined by the marker panels (**Fig. 4f,4e**). In comparison, Tangram, cell2location and CARD offered approximations of the cell-type distribution, while Seurat and SpatialDWLS produced results dominated by oligodendrocytes and neurons. On the other hand, Stereoscope, DestVI, and SPOTlight exhibited notably inferior deconvolution performance when applied to MS-based spatial proteomics data. Notably, while Stereoscope is capable of identifying B cells enriched in follicles in human palatine tonsil data derived from ADTs (**Fig. 3f**), it is unable to replicate similarly clear results when applied to MS-based mouse brain coronal data (**Fig. 4f-g**). This highlights the substantial impact of assumptions regarding data distribution on the analysis of spatial proteomics data using deconvolution methods. In summary, our results conclusively established that Spatial-DC exhibited superior performance for MS-based spatial proteomics deconvolution tasks.

Next, we analyzed the reconstructed spatially and cell-type resolved and proteomic profiles of the mouse coronal brain data by Spatial-DC (**Fig. 4h,i; Fig. S9b and Table S6**). Comparison of differential expression proteins (DEPs) between the original spatial proteomics data and the reconstructed spatially and cell-type proteomics data demonstrated that the resolution of the spatial proteomics data was improved and the cell-type-specific information, originally mixed within the spatial spots, became clearly distinguishable after deconvolution by Spatial-DC (**Fig. 4k; Methods**). Gene Ontology Biological Processes (GO-BP) enrichment analysis was performed to validate the cellular functions associated with these reconstructed proteomic profiles (**Fig. 4k; Methods**). We then studied cell-cell communications with the reconstructed proteomic profiles of the coronal section. Among the signaling pathways present in the data, the signaling protein complex involving neural cell adhesion molecule (NCAM) and *EphB/ephrin B* (EPHB) exhibited a clear spatial orientation during the contact of coronal cells (**Fig. 4l,m; Table S6**). In the nervous system, NCAM has been demonstrated to play a pivotal role in mediating cell adhesion. The binding of these molecules is crucial in influencing various neuronal developmental processes (i.e., cell migration, proliferation and regeneration)^53,54^. Our analysis of cell-cell communications within neurons elucidated the directional flow of the NCAM signaling pathway, originating from the CTX region and transmitting to the HIP, TH and HY regions, aligning with the primary roles and functions of NCAM signaling in the nervous system (**Fig. 4a,l**)^53,54^. We also conducted an analysis of the EPHB signaling pathway, which is involved in the regulation of nerve injury through interaction between neurons and glial cells^55^. Our findings highlighted a distinctive aspect of intercellular communication between neurons and astrocytes, mediated through the EPHB signaling pathway. Notably, we observed a prominent enrichment of signals transmitted via this pathway, specifically in the HIP and TH regions. This observation indicates the regional occurrence of EPHB signaling between these two cell types, emphasizing the importance of spatial context in regulating intercellular signaling and communication within the coronal plane of the brain. (**Fig. 4a,m**).

These results clearly showcased the deconvolution capability of Spatial-DC on spatial proteomics data of spatially-resolved MS technology. Comprehensive evaluations on publicly available spatial proteomics datasets generated by two major technological categories suggested that Spatial-DC is broadly applicable to a diverse range of spatial proteomics data, serving as a robust and valuable tool for analyzing and interpreting spatial proteomic information.

### Application of Spatial-DC for tumor microenvironment (TME) analysis of pancreatic cancer on self-collected dataset

In evaluating the deconvolution performance on real-world datasets of Spatial-DC and other methods for certain cell-types, in the absence of definitive ground truth, we compared the cell-type distribution with the abundance of their respective signature proteins in each spatial spot. However, it is important to acknowledge that the accuracy of these evaluations may be compromised by limitations in acquisition technologies and the potential interference from highly abundant proteins, resulting in incomplete quantitative assessments. Moreover, given the intricate and vastly variable nature of the TME, it is imperative to undertake a comprehensive systemic assessment of the deconvoluted spatial proteomics data within this disease environment. Consequently, we acquired self-collected spatial proteomics data from a genetically engineered mouse model of pancreatic ductal adenocarcinoma (PDAC), utilizing laser microdissection (LMD) coupled with by LC-MS/MS for detection (**Fig. 5a; Methods; Table S3 and Note S1**). The spatial proteomics data consists of 108 sample slices with 100-μm spot diameters and includes a total of 3,607 identified proteins. Each sample contains an average of over 1,500 proteins identified, highlighting the rich proteomic landscape captured (**Fig. S10a,S10b**). To unravel the intricacies of this complex TME, we collected corresponding single-cell-type proteomics data of five PDAC mice acquired via FACS sorting followed by LC-MS/MS^56^. This reference single-cell-type proteomics data comprises 50 samples and 5,788 identified proteins, spanning three distinct cell lineages: pancreatic cancer cells (PC), pancreatic cancer-associated fibroblasts (CAF), and pancreatic immune cells (PI). Among the PI, there are four subtypes of lymphoid cells and four subtypes of myeloid cells. The lymphoid cell subtypes include CD4 T cells (T4), CD8 T cells (T8), regulatory T cells (Treg) and B cells (B), while the myeloid cell subtypes consist of neutrophils (NEU), macrophages (MAC), monocytes (MO), and dendritic cells (DC). We then conducted UMAP analysis, which revealed a distinct separation among the ten cell types, enabling clear differentiation between PC, CAF, and PI (**Fig. 5b**). Upon integrating the reference proteomics and spatial proteomics data, we identified 2,854 proteins common to both datasets (**Fig. S10c**), providing a valuable overlap for further investigation.

**Fig. 5.**
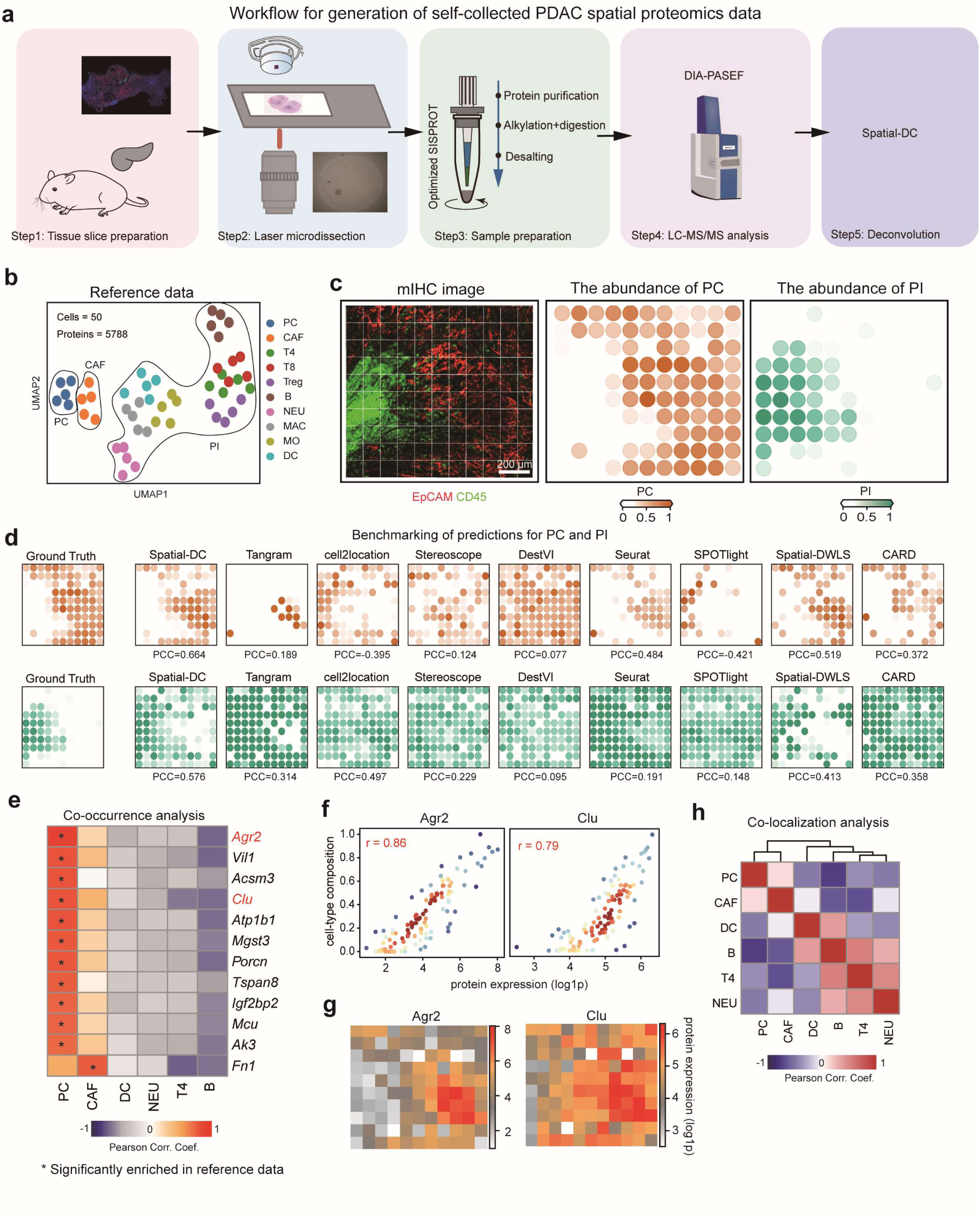
Application of Spatial-DC for cell-type distributions on self-collected mouse PDAC data. **a-c,** Overview of the self-collected spatial proteomics data. **a,** The pipeline outlines the major steps involved in the generation of self-collected spatial proteomics data, encompassing the preparation of the tissue slice from KPC mouse, acquisition of target spatial proteomics using laser microdissection (LMD)-based LC-MS/MS, determination of the related cell-type abundance through mouse tissue mIHC (**Methods**), and conduction of deconvolution by Spatial-DC. **b,** UMAP visualization of the reference single-cell-type proteomics^56^ derived from KPC mice. **c,** Visualization of the spatial co-localization of pancreatic cancer cells (PC) and pancreatic immune cells (PI) in a merged two-plex antibodies mIHC image (left. The abundance of PC and PI is represented by the fluorescence strength of *EpCAM* and *CD45*, respectively (middle and right) (**Methods**). This serves as the ground truth for the distribution of PC and PI. The color scale in the plot ranges from 0 to 1, indicating the scaled abundance of cells, where darker colors represent higher abundance. **d,** Spatial maps of cell-type distributions of PC (upper row) and PI (bottom row) for the ground truth and the predictions of Spatial-DC and eight competing methods, with the PCC metrics between the ground truth and the corresponding prediction indicated on below (**Methods**). **e-f,** Applications of cell-type composition predicted by Spatial-DC. **e,** Heat map displaying the Pearson correlation coefficient between the predicted cell-type composition and the original protein expression across 108 spatial spots. Protein names are shown in the right side of the heat map, with those of two representative PC-correlated proteins colored red. Asterisks represents proteins significantly enriched in reference data (**Fig. S10g; Methods**). **f,** Scatter plots showing the linear correlation between log1p-normalized protein abundance from spatial proteomics data and predicted cell-type composition of PC by Spatial-DC for two representative PC-correlated protein, *Arg2* and *Clu*, respectively (**Methods**). The PCC metrics are labeled on the graphs. **g,** Spatial maps depicting the log1p-normalized abundance of representative PC-correlated proteins corresponding to **f**. **h,** Unsupervised clustering analysis of Pearson correlation coefficient using predicted cell-type composition by Spatial-DC, indicating the co-localization of the six cell types (**Methods**).

To assess the deconvolution performance of Spatial-DC, we designed a two-plex antibody panel to quantify the abundance of major cells in the tissue slice of spatial proteomics data. The antibody panel comprised *EpCAM* for visualizing PC and *CD45* for PI (**Fig. S10d; Methods**). The image channels corresponding to PC and PI were individually transformed into the ground truth representation of their respective cell-type distributions (**Fig. 5c; Methods; Table S7**). Then, we performed a comprehensive evaluation analysis on the mouse PDAC data. Upon visualizing the outcomes, we found that Spatial-DC delivered the most accurate cell-type distribution compared to the ground truth. While Tangram, CARD, SpatialDWLS and Seurat also produced reasonable predictions, their performance fell short of Spatial-DC. In contrast, cell2location, Stereoscope and DestVI significantly underperformed in this experiment (**Fig. 5d**). The quantitative assessment further corroborated the conclusion, revealing that Spatial-DC surpassed all other comparative methods across various evaluation metrics (**Fig. 5d**), showing PCC metrics of 0.664 and 0.576, and SPCC metrics of 0.696 and 0.530 for the PC and PI, respectively (**Fig. 5d; Table S7**).

Assuming that the abundance of cell-type-specific markers is directly proportional to the composition of specific cell types within a sample, we aimed to uncover potential cell-type-specific proteins quantified from the original spatial proteomics data. To achieve this, we employed a co-occurrence analysis approach using the predicted cell-type composition by Spatial-DC, which allowed us to identify proteins that consistently occur in association with particular cell types, providing insights into their potential role as cell-type-specific markers at spatial level (**Methods**). Our comprehensive analysis identified a multitude of proteins that show promise as cell-type spatially correlated and cell-type-specific markers for both the spatial proteomics data and the reference data (**Fig. 5e; Fig. S10g and Table S7**). For instance, anterior gradient protein 2 homolog (*Agr2*) and clusterin (*Clu*) emerged as prominent candidates for PC, as evidenced by their distinct patterns of expression (**Fig. 5f-g**). These findings suggested that *Agr2* and *Clu*, along with other identified proteins, may hold significant potential for accurately identifying and characterizing different cell types within complex biological samples. In addition, our co-localization analysis uncovered intriguing relationships among different cell types (**Methods and Table S7**). Specifically, we observed that PC and CAF exhibited closer proximity to each other, potentially indicating functional interactions or shared microenvironments. On the other hand, DC and B cells displayed a high degree of similarity, which may be attributed to their shared role in antigen presentation (**Fig. 5h**). The analysis of spatial domains further validated the effectiveness of our Spatial-DC approach in deconvolving cell-type composition (**Fig. S10e-f**). Our findings demonstrated that the regions identified by Spatial-DC were more accurately aligned with the patterns observed in mIHC compared to the clustering results based on original spatial proteomics data (**Fig. S10d-f**).

In summary, our findings conclusively demonstrated the superiority of Spatial-DC over competing deconvolution methods when applied to our self-collected spatial proteomics data of pancreatic cancer. The comprehensive analysis not only underscored the reliability and precision of our approach but also provided valuable biological insights into the intricate cellular landscapes present within complex disease tissue slices.

### Construction of protein-protein interaction (PPI) network of pancreatic cancer at cell-type and spatial levels from proteomic profiles reconstructed by Spatial-DC

Next, we utilized Spatial-DC to analyze the reconstruction outcomes of cell-type proteomic profiles derived from mouse PDAC data by Spatial-DC (**Fig. S10g**). Our findings revealed that over 2,500 proteins were successfully reconstructed, encompassing 582 samples across six distinct cell types (**Fig. S10h; Table S8**). These cell types exhibited clear specificity in the UMAP plot (**Fig. 6a**). Upon performing statistical analysis on the reconstructed data (**Methods**), we identified several well-established cell-type-specific marker proteins within these profiles. For instance, PC were characterized by markers such as *Cdh1*, *Krt19*, and *Epcam*, CAF were identified by *Vim*, *Acta2*, and *Dcn*, and NEU cells were distinguished by *S100a9*, *Mpo*, and *Elane* (**Fig. 6b and Table S10**). GO-BP enrichment analysis corroborated that the enriched terms aligned with the specific functional roles of the corresponding cell types. For example, the functional categories “mitochondrial translation” and “cell-cell adhesion mediated by cadherin” were enriched in PC, reflecting their metabolic and structural roles. Similarly, CAF showed enrichment in “extracellular matrix organization” and “collagen fibril organization,” indicating their crucial role in the remodeling of the tumor microenvironment. These statistical and functional enrichment analyses reinforced the precision and reliability of Spatial-DC in reconstructing cell-type-specific proteomic profiles and capturing the fundamental cellular functions (**Fig. 6c and Table S8**).

**Fig. 6.**
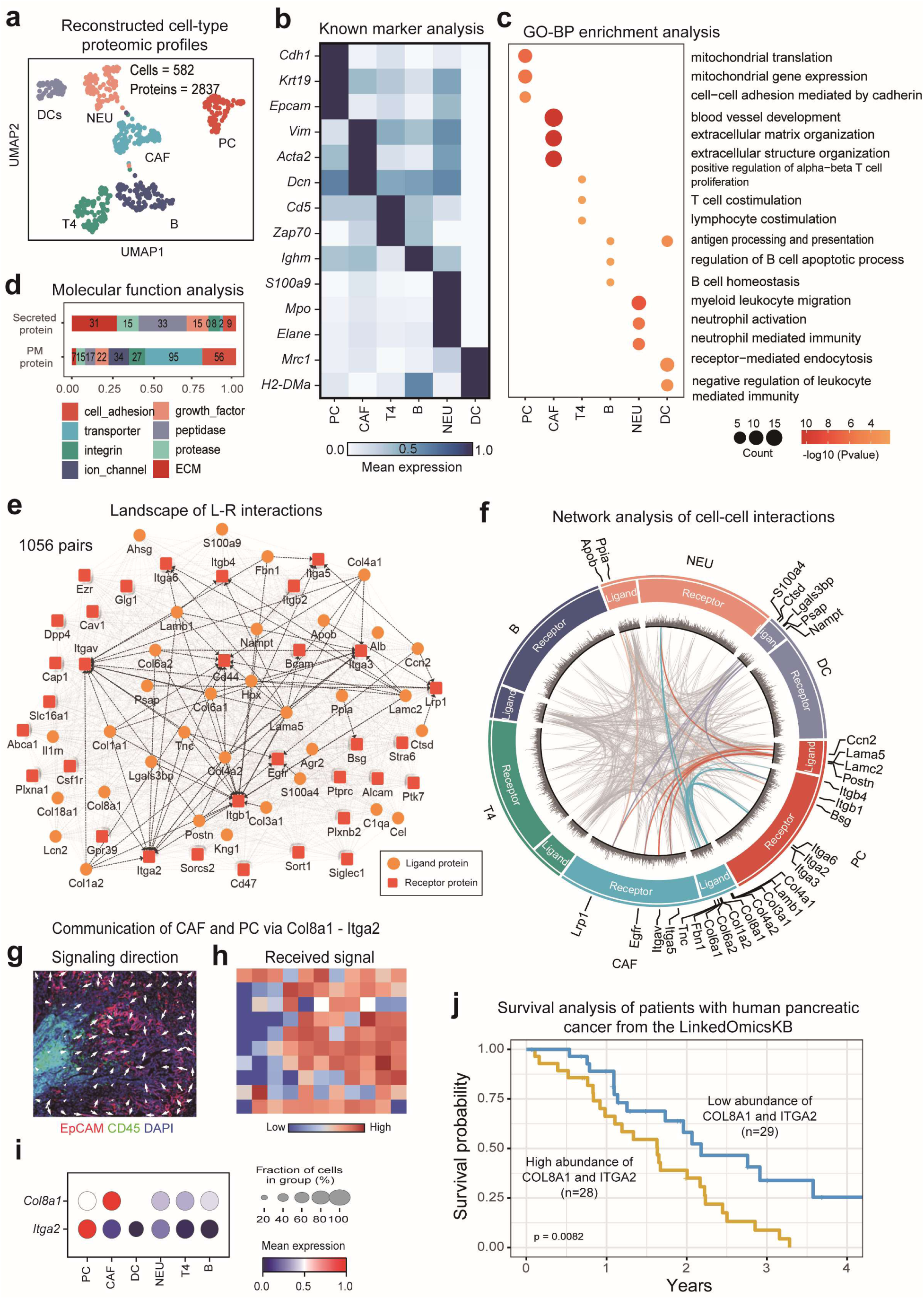
Application of Spatial-DC for proteomic profiles and cell-cell communication on self-collected mouse PDAC data. **a-d**, Functional analysis of spatially and cell-type resolved proteomic profiles reconstructed by Spatial-DC. **a,** UMAP of six reconstructed proteomic profiles (**Methods**). **b,** Heat map showing the mean abundance of representative well-known cell-type-specific markers (**Methods**). **c,** Dot plot exhibiting 18 significantly enriched GO-BP terms for the six cell types using cell-type-specific proteins. **d,** Stacked bar chart illustrating the number of quantified secreted proteins and plasma membrane (PM) proteins within eight major categories across the six distinct cell types from reconstructed cell-type proteomic profiles. The counts of proteins within each category are indicated on the respective segments of the bars. **e-j**, Construction of cellular crosstalk using reconstructed proteomic profiles. **e,** Visualization of identified L-R interactions using reconstructed secreted proteins and PM proteins (**Methods**). Ligand proteins are colored orange, while receptor proteins are colored red. Known L-R interactions are linked with black dash line. **f,** Circos plot illustrating cell-cell interactions among the six cell types using reconstructed proteomic profiles (**Table S8**). The names of cell types are labels on the outermost ring. The first inner layer is segmented into twelve sections, with each cell type comprising two sections for its ligands and receptors. The cell-type-specific ligand or receptor proteins are labeled. The second layer, presented as bars, illustrates the intensity of the related proteins. The innermost layer visualizes the lines connecting the interacting proteins, with those of cell-type-specific ligand proteins colored as same as sender cells. **g-h**, Representative cell-cell communication for mouse PDAC data between CAF and PC via *Col8a1* and *Itga2*. The signaling direction and the strength of received signal are represented in **g** and **h**, respectively (**Methods**). **i**, Dot plot demonstrating the mean expression of *Col8a1* and *Itga2* from reconstructed cell-type proteomic profiles by Spatial-DC, the size of dot representing the fraction of cells of reconstructed spots. **j**, Survival analysis on pancreatic patients from LinkedOmicsKB pancreatic data^38^, depicting the relationship between interaction score and patient survival time (by years). The pancreatic patients are grouped into two categories, low-risk and high-risk, based on the co-expression levels between *COL8A1* and *ITGA2* (**Methods**).

Analysis of Gene Ontology Molecular Function (GO-MF) indicated that Spatial-DC effectively reconstructed essential functional proteins, such as growth factor, cell adhesion molecules and extracellular matrix (ECM) proteins (**Fig. 6d**). Notably, these proteins are either anchored to the plasma membrane or secreted into the extracellular space, emphasizing their pivotal role in orchestrating intercellular signaling cascades through intricate cellular crosstalk within the tumor microenvironment of pancreatic cancer^57^. To explore cellular crosstalk mediated with such proteins in pancreatic cancer, we manually curated a mouse ligand-receptor (L-R) interactions database using the reconstructed secreted proteins and plasma membrane proteins (**Methods and Table S8**). The database we assembled could serve as a pivotal resource for delving into the complex molecular interactions that propel the progression of pancreatic cancer. Our reconstructed spatially and cell-type resolved proteomics data encompasses 1,056 ligand-receptor protein interactions derived from 33 secreted proteins and 32 plasma membrane proteins. Notably, among these interactions, 77 L-R interactions involving 23 secreted proteins and 12 plasma membrane proteins have been confirmed as known interactions (**Fig. 6e, Table S8**).

Subsequently, we employed a statistical strategy to analyze the sender and receiver cells of the 1,056 L-R interactions (**Methods**). Our analysis pinpointed 128 ligand proteins and 66 receptor proteins originating from the six distinct cell types, resulting in the quantification of 1,980 cell-cell L-R interactions. Notably, 49 of these interactions emerged as statistically significant and were predominantly observed among PC, CAF, NEU, and DC (**Fig. 6f; Methods; Table S8**). This finding highlighted the specific and dynamic nature of cellular crosstalk within the TME of pancreatic cancer. Of particular interest, collagen type VIII alpha-1chain (*Col8a1*) and integrin alpha-2 (*Itga2*) stood our as significant L-R interaction in the reconstructed spatially and cell-type proteomic profiles (**Fig. 6f,i**). While the interplay between *Col8a1* and *Itga2* have been documented^58^, its pivotal role in mediating the intricate intercellular communication between PC and CAF in pancreatic cancer remains largely unexplored. Our observations led us to propose that the paracrine signaling pathway mediated by *Col8a1*-*Itga2* L-R interaction operates within the pancreatic cancer microenvironment (**Fig.6g-i; Methods**). Our detailed analysis of cell-cell communication between CAF and PC, specifically mediated by *Col8a1-Itga2*, elucidated the signaling direction and primary flow, which was predominantly observed in non-immune regions (**Fig. 6g,h**). This finding is further supported by our mIHC results, indicating that the signaling is enriched in non-immune areas (**Fig. 6g,h**). The close proximity of CAF and PC, coupled with their significantly elevated expression of ligand and receptor proteins, facilitated these vital communication (**Fig. 6g,h,i; Methods**). Moreover, our analysis of survival curves in pancreatic cancer patients implied a notable impact on prognosis of high co-expression scores of *COL8A1*-*ITGA2* compared to those with low expression scores (**Fig. 6j; Methods; Table S8**). This observation underscored the clinical significance of this L-R interaction in pancreatic cancer, suggesting that it may serve as a prognostic marker or potential therapeutic target. The high co-expression of *COL8A1* and *ITGA2* may reflect an active crosstalk between cancers cells and surrounding microenvironment, which contributes to the aggressive nature of pancreatic cancer and impacts patient survival.

In summary, our study underscored the transformative potential of Spatial-DC in systematically unraveling complex cell-cell interactions and signaling direction within spatial contexts, leveraging self-collected spatial proteomics data. This approach offers profound biological insights into the intricate tumor microenvironment, providing a novel lens to dissect its complexities and potentially inform improved patient outcomes. By elucidating critical ligand-receptor interactions such as *COL8A1*-*ITGA2*, our findings pave the way for future research targeting these molecular mechanisms to develop more effective therapies for pancreatic cancer and other malignancies.

## Discussion

In recent research, various deconvolution methods have emerged in the field of spatial transcriptomics. However, these methods are primarily designed based on the distribution characteristics of transcriptomic data and face limitations when directly applied to proteomics data. Given the advantages of deep learning tools in capturing complex implicit relationships in representation learning, we propose Spatial-DC to leverage these tools to learn the inherent association between proteomic expression and cell-type resolved information and then incorporate the intrinsic characteristics of the spatial proteomics data itself through a self-supervised learning strategy. Spatial-DC predicts cell-type composition as well as reconstructs spatially and cell-type resolved proteomic profiles for spatial proteomics data through two deep learning techniques in two stages, namely transfer learning and self-supervised learning. Transfer learning involves capturing cell-type-resolved information from the reference proteomics data and transferring it to the spatial proteomics data to conduct an intermediate prediction for each spot. The intermediate prediction inevitably introduces bias due to the significant gap between reference data and spatial proteomics data. The GCN with self-supervised learning strategy is then utilized to address this bias and integrate spatial information by incorporating the intermediate prediction of cell-type-resolved information with the characteristic of spatial proteomics data, including spatially neighborhood information of spots and corresponding proteomic profiles, to obtain the final refined cell-type-resolved information prediction (**Methods and Fig. S6**).

To validate the feasibility and performance of Spatial-DC in the deconvolution of spatial proteomics, we compared it with eight state-of-the-art methods on both synthetic spatial proteomics data and real-world datasets derived from various tissues through different spatial proteomics technologies. The experiment results underscored that Spatial-DC accurately and robustly predicted cell-type composition and provided the reconstructions of spatially and cell-type resolved proteomic profiles with high congruence to biological markers and characteristics. In our experiments, Spatial-DC demonstrated performance that surpassed the competitive methods across all the datasets, showcasing its superiority and generalizability in spatial proteomics data.

Furthermore, we collected a spatial proteomics dataset from pancreatic cancer mouse to demonstrate the application and subsequent downstream analyses of Spatial-DC. By utilizing cell-type composition deconvoluted by Spatial-DC, we were able to identify specific proteins that demonstrated reliability as spatially correlated and cell-type-specific markers for both spatial proteomics data and reference data (i.e., *Agr2* and *Clu* for PC) (**Fig. 5f; Fig. S10g**). These findings offer a fresh perspective, suggesting that the predicted cell-type composition holds considerable promise for accurately identifying and characterizing cell-type-specific markers in both spatial and reference proteomics data. Besides accurately predicting cell-type composition compared to the ground truth derived from mIHC experiments, Spatial-DC can also provide robust and meaningful reconstructions of spatially and cell-type resolved proteomic profiles, validated by GO-BP and GO-MF analyses. The reliability of the reconstructed proteomic profiles by Spatial-DC ensures the ability to conduct in-depth studies of cell-cell communication based on L-R interaction. In this work, we used the interaction between *COL8A1* from CAF and *ITGA2* from PC as an example, which was discovered in the TME of pancreatic cancer and significantly associated with the prognosis of pancreatic cancer patients, showing the great potential of Spatial-DC to uncover new clinical findings.

Although the excellent performance of Spatial-DC has been proven in various spatial proteomics data, there are still some areas that require improvement. One primary limitation of Spatial-DC is its reliance on reference single-cell or single-cell-type proteomic profiles. The reference proteomics data, as the sole source of cell-type resolved information, is crucial throughout the entire deconvolution process. It is imperative to underscore the pivotal role that the quality of reference proteomics data and the discrepancy between reference and target spatial proteomics data play in influencing the efficacy of deconvolution methods (**Fig. 2h; Fig. S3-S5**). Almost all deconvolution methods, including Spatial-DC, are constrained by this limitation. Spatial-DC attempts to alleviate the impact of such discrepancy by employing a two-stage approach with a self-supervised learning strategy in the second stage. Nonetheless, the quality of reference proteomics data remains a crucial factor that can still influence the performance of our proposed method. Consequently, future research could explore alternative methodologies to optimize the utilization of reference proteomics data, such as introducing more reliable normalization and batch correction procedures in pre-processing or applying more robust transfer learning techniques. If data discrepancies can be effectively alleviated, data from other modalities could be used as reference data, such as the abundant single-cell RNA sequencing datasets, yielding valuable insights^59^. Moreover, the growing prevalence of large-scale pre-trained models offer a new direction to potentially address this issue by integrating extensive information about the implicit correlation between protein expression and cell types from different sources during the pre-training phase, thus leading to reference-free methods for the advanced deconvolution of spatial proteomics data^60,61^.

Another limitation of Spatial-DC is that the reconstructed proteins are confined to the intersection of commonly identified proteins between the reference proteomics data and the target spatial proteomics data. However, some meaningful proteins are identified only in either the spatial proteomics data or the reference proteomics data due to limitations in detection sensitivity or quantitative strategies. Note that this challenge is also linked to the utilization of reference data and will be tackled through innovative use of reference data or the advent of reference-free methods. Before that, investigating appropriate imputation methods based on the characteristics of each dataset (e.g., the proportion of missing values^62^) will be one of our future directions.

In summary, we proposed Spatial-DC, a versatile and powerful tool for predicting cell-type composition and reconstructing spatially and cell-type resolved proteomic profiles from spatial proteomics data. The effectiveness and potential of Spatial-DC were evaluated through various experiments on diverse tissues from different technologies. In fact, Spatial-DC is not bound by the resolution of spatial proteomics data. With the continuous advancement of spatial proteomics techniques, resulting in increasingly higher resolutions of capturing spots, Spatial-DC remains applicable and can ultimately be employed to perform cell type annotation for spatial proteomics when reaching single-cell or even sub-cellular resolution^31^. Overall, we anticipate that Spatial-DC will provide valuable insights into dissecting the complex interplay between cells in tissue microenvironments through spatial proteomics, facilitating a more comprehensive understanding of disease biology and identification of novel therapeutic targets.

## Methods

### Overview of Spatial-DC

#### Introduction

Spatial proteomics faces a significant challenge in deconvoluting mixed spots, which often represent contributions from multiple cell types. Spatial-DC addresses this issue by integrating spatial proteomics data (spatial data) with relevant cell-type resolved information derived from reference single-cell or single-cell-type proteomics datasets (reference data). This approach enables Spatial-DC to estimate both the cell-type composition and corresponding cell-type proteomic profiles within these mixed spots. Spatial-DC is designed to exploit the underlying relationship between the spatial data and the cell-type resolved information informed by reference data.

To construct Spatial-DC, we first train a distribution model **DIS** of deep neural network (DNN). This model is trained using synthetic spots data generated from the reference data, enabling it to learn the complex connections between proteomics data and cell-type resolved information. Once trained, the model is applied to spatial proteomics data to obtain intermediate prediction for cell-type resolved information from the spatial data.

Recognizing the importance of spatial distance in capturing dependencies between cells, as nearby cells tend to have similar or related expression patterns. To incorporate this information, as well as the characteristics of spatial proteomic profiles, we build a self-supervised model using a graph convolutional network (GCN), denoted as model **SSM**. This model leverages the proteomic profiles and spatial coordinates of each spot to refine the intermediate prediction of cell-type resolved information, resulting in refined cell-type resolved information based on characteristics of the spatial data, which are subsequently converted to cell-type composition as well as cell-type resolved proteomic profiles for each spot.

In the subsequent section, we present a comprehensive overview of the core process for constructing Spatial-DC, including the generation of training data, the structure of the distribution model **DIS** and the self-supervised model **SSM**, and the process for obtaining cell-type composition and cell-type proteomic profiles. The details of the model structure, involving the hyperparameters settings and model components, are listed in **Fig. S1 and Table S1**.

#### Generation of training data

To train the distribution model **DIS**, we leverage a set of reference data to generate synthetic spots data. Let an ***c*** × ***p*** matrix ***R*** represents the reference proteomics data with ***c*** cell types and ***p*** proteins. We first use the ‘random.dirichlet’ function of the NumPy package to obtain synthetic cell-type composition matrix ***Q*** with ***c*** cell types and ***m*** spots sampled from a Dirichlet distribution. Let ***Q****_ij_* donate the *j*-th cell-type composition within *i*-th synthetic spot, which is expressed as:

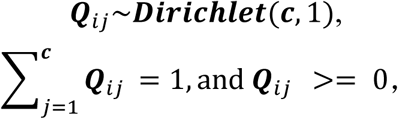

Let an ***m*** × ***p*** matrix ***M*** donates the expression profiles of synthetic spots with ***m*** spots and ***p*** proteins.

Then we got:

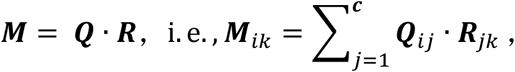

***M***_ik_ represents the *k*-th mixed protein expression within the *i*-th spot. In this formulation, we assumed that there are representative proteomic profiles for each cell type to conduct simulations. If the cell-types in the reference data include more than one replicate, random sampling would be applied to these replicates to incorporate variance into the construction of the training data. Let an ***m*** × ***c*** × ***p*** matrix ***N*** represents the weights of expression profiles for each cell type in synthetic spots, which is expressed as:

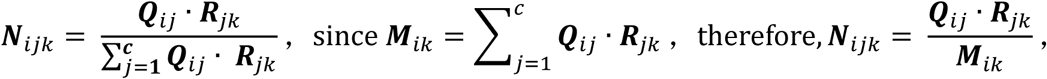

***N***_ijk_ represents the contribution of the *k*-th protein expression of the *j*-th cell type within the *i*-th spot to the *k*-th protein expression of the entire spot. Therefore, for all *j*, it holds that 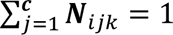. Our proposed approach utilizes synthetic spots ***M***, synthetic cell-type composition ***Q***, and synthetic cell-type proteomic weights ***N*** to train the **DIS** model. Specifically, ***M***, with dimensions of ***m*** × ***p***, serves as the input to the model. Meanwhile, ***Q***, with dimensions of ***m*** × ***c***, and ***N***, with dimensions of ***m*** × ***c*** × ***p***, are concatenated to form synthetic labels ***L***, with dimensions of ***m*** × ***c*** × (1*+**p***), which serves as the training labels for the **DIS** model.

#### Structure of the distribution model

The distribution model **DIS** is a DNN composed of three layers, each comprising a dropout layer, a linear layer, and a non-linear activation function. The activation function utilized in the first two hidden layers is CELU, while ReLU is employed in the output layer.

Dropout regularization is used to improve the generalization ability. We use the synthetic spots data ***M*** as the input and the synthetic labels ***L*** as the training labels to train **DIS**. Mean Squared Error (MSE) is employed as the loss function, and Adam optimizer is used to optimize the parameters with backpropagation.

To reduce the various effects (e.g. cross-platform and batch-effect), we utilize the same preprocessing approach to prepare the input data. We perform log1p operation to the intensity of proteins for the synthetic spots ***M*** and the spatial proteomic profiles ***S_EXP_***. Then, we use the ‘scalè function from the sklearn package to scale both matrices to have a mean of zero and a standard deviation of one.

#### Structure of the self-supervised model

The self-supervised model, denoted as **SSM**, comprises three main components: a deep autoencoder (**DAE**), a variational graph autoencoder (**VGAE**), and a classifier (**CLS**). **DAE**, including an encoder (**D-E**) and a decoder (**D-D**), is used to learn the compressed representation of the spatial proteomic profiles ***Sexp*** of the spatial proteomics data. **D-E** is a DNN with two layers, each comprising a dropout layer, a linear layer and CELU activation function. **D-D** comprises a linear layer. **VGAE**, composed of a GCN^63^ encoder (**G-E**) and an inner product decoder (**G-D**), is used to learn the latent graph-structured representations of spatial proteomics data. **CLS** is a DNN with a linear layer and a ReLU activation function that takes the concatenated embedding ***Z*** and produces the final prediction. The total loss ***Loss***_GCN_of the self-supervised model comprises three components: Firstly, there is the reconstruction loss of **VGAE *Loss***_VGAE_that aims to reconstruct the adjacency matrix derived from the transformation of ***Scoo***. Secondly, **DAE** contributes to its own reconstruction loss ***Loss***_DAE_, which focuses on reconstructing ***Sexp***. Lastly, the cross-entropy loss ***Loss***_CLS_is incurred by aligning the outputs of **CLS** with the intermediate prediction ***P***, ensuring that the classification results are coherent with the encoded information within ***P***. Noted that the intermediate prediction ***P*** is derived from the **DIS** model with the spatial proteomic profiles ***Sexp*** from the spatial proteomics data as the input. The overall loss function is formulated as follows:

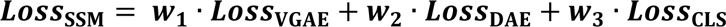

where ***w_1_***, ***w_2_*** and ***w_3_*** represent the weights assigned to each of the three loss components, respectively.

The self-supervised model focuses on representing spatial proteomics data as undirected graphs and is capable of learning interpretable latent representations. **D-E** takes the spatial proteomic profiles ***Sexp*** as input and converts it to a lower-dimensional embedding ***Z1***, which is then taken by **G-E** as the feature matrix. **G-E** takes the graph-structured data, which is represented by an adjacency matrix ***A*** and the feature matrix ***Z1***, as input and converts it to the embedding ***Z2***. To achieve ***A***, Euclidean distance of k-nearest neighboring spots is calculated using the spatial coordinates ***Scoo*** from the spatial proteomics data. ***Z1*** and ***Z2*** are concatenated to obtain the spatial embedding ***Z***. **D-D** is used to convert ***Z*** to the reconstructed proteomic profiles 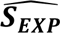, **G-D** to convert ***Z*** to the reconstructed adjacency matrix 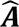, and ***CLS*** to convert ***Z*** to the refined cell-type prediction ***Y***. The architecture of **SSM** is depicted in Fig. S1.

#### Obtain cell-type composition and proteomic profiles

The refined cell-type prediction ***Y***, with dimensions of ***n*** × ***c*** × (1*+**p***), akin to the intermediate prediction ***P*** and the synthetic labels ***L*** in structure, is then split along the third dimension, yielding two components: cell-type composition ***D***, an ***n*** × ***c*** dimensional matrix representing the composition of cell types in the spatial proteomics data (deconvolution results), and cell-type proteomic profile weights ***W***, an ***n*** × ***c*** × ***p*** dimensional matrix capturing the weights of proteomic profiles for each cell type. ***n*** denotes the number of spots in the spatial proteomics data. Given the proteomic profile weights ***W*** and the spatial proteomic profiles ***S_EXP_*** (***n*** × ***p*** dimensional, abbreviated as ***S*** in subsequent equations) from the spatial proteomics data, the cell-type proteomic profiles ***E***, representing the reconstructed proteomic profiles for each cell type, can be derived with:

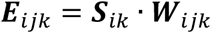

***E****_ijk_* represents the *k*-th protein expression of the *j*-th cell type within the *i*-th spot. With this, the entire process of deconvolving the target spatial proteomics data and reconstructing the cell-type proteomic profiles for individual cell types is completed. The final outputs of Spatial-DC are the cell-type composition ***D***, with dimensions of ***n*** × ***c***, and the reconstructed cell-type proteomic profiles ***E***, with dimensions of ***n*** × ***c*** × ***p***.

For one spot containing various cell-types with un-equal cell counts, to compare the cell-type proteomic profiles in the following statistical analysis, we use cell-type composition ***D*** to normalize ***E***. Let ***E***’ donates normalized cell-type proteomic profiles, then we have:

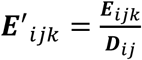

### Generation of synthetic data

We obtained 36 synthetic spatial proteomics datasets and 12 synthetic reference proteomics data (**Table S2**). These spatial datasets incorporated spatial localization information derived from well-defined, single-cell resolved structures. For the synthetic spatial proteomics data, we varied the spot sizes (100, 150, and 200 μm) and introduced differing levels of noise (25%, 50%, and 75%) as well as varying numbers of variable proteins (ranging from 800 to 200 top and bottom variable proteins) to simulate a range of conditions in spatial proteomic data. Additionally, for the reference single-cell-type proteomics data, we applied the same levels of noise and same panel of variable proteins to perform simulations.

#### Obtain single-cell resolved tissue structure

To accurately mimic the complex tissue structure observed in spatial proteomics data from real-world samples, we employed the precisely annotated spatially and single-cell resolved NSCLC from the CoxMx SMI platform (www.nanostring.com), which are publicly accessible at https://nanostring.com/resources/smi-ffpe-dataset-lung13-data. These tissue structure data provide precise locations for cells, enabling the generation of synthetic spatially and single-cell resolved data that reflects the context of naïve tissue microenvironment.

#### Obtain synthetic spatially and single-cell resolved NSCLC data with well-defined tissue structure

We generated synthetic spatially and single-cell resolved data by utilizing single-cell proteomics data as the basis for projection. This projection data was then used to replace the cells within a well-defined tissue structure through a sampling process. Specifically, we maintained the commonly identified cell types in both tissue structure and the projection data. For each cell type within the tissue structure, we fixed the locations of the cells and performed a random sampling from the projection data to fill in the proteomic profiles of the cell types at their respective spatial locations. In the case of the synthetic NSCLS spatially and single-cell resolved data, we substituted the cell locations within the tissue structure with a single-cell proteomics dataset that comprised three distinct cell types, totally 32,859 cells, and 1,003 proteins.

#### Generate synthetic spatial proteomics datasets with varying spot sizes and external noise

To systematically explore the effects of various factors on synthetic spatial proteomics data, we developed a simulation approach that incorporates two key conditions on synthetic data: spot-level variations in cell density and the introduction of external noise. To simulate variations in cell density at the spot level, we created synthetic spots with different sizes (100, 150 and 200 μm), where each spot contains a varying number of cells, ranging from one to a hundred. The proteomic profiles of each synthetic spot were generated by summing the proteomic abundances of the cells within a simulated grid, with the coordinates of the spot centered on the grid. To introduce external noise into synthetic spot-level datasets, we first constructed a Gaussian distribution based on the spot-level proteomic profiles. We then sampled from this distribution at varying degrees (25%, 50%, and 75%) and added it to the original proteomic profiles. The external noise added to the reference data was consistent with the spatial data.

### Performance evaluation

In this study, we performed the deconvolution on synthetic, public available and self-collected spatial proteomics datasets. To evaluate the ability of deconvolution, we used four metrics, including PCC, SPCC, CCC and RMSE, to assess Spatial-DC and competing methods.

The following equations were used to calculate these metrics, respectively:

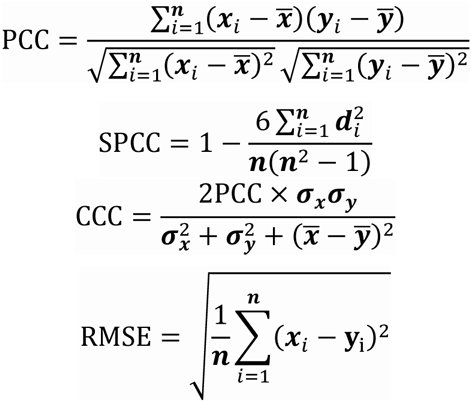

where ***n*** is the number of spots, ***x***_i_ and ***y***_i_ represent the cell-type composition of the *i*-th spot in the ground truth and the predicted results, respectively, 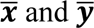 are the mean values of ***x*** and ***y***, ***d****_i_* is the difference in ranks between ***x***_i_ and ***y***_i_, ***σ***_***x***_ and ***σ***_***y***_ represent the standard deviations of ***x*** and ***y***.

In addition, the four metrics were calculated for all cell types from spots of datasets used in the synthetic NSCLC data. To evaluate the cell-type composition in a same range, the total composition of all cell types was normalized to 1 in each spot.

For public datasets lacking direct ground truth for cell-type composition in each spot, we utilized the expression of known cell-type-specific markers (termed “signature scores” in this studies) from original spatial proteomics data as an indirect guidance. This approach serves as a valuable surrogate metric for the evaluation of deconvolution algorithm^14,23^. For the mouse brain coronal data, we chosen three cell-type-specific marker panels to calculate signature scores: three proteins (*Cacng3*, *Slc30a3*, *Synpo*) for neurons, five proteins for oligodendrocytes (*Mbp*, *Plp1*, *Cnp*, *Cldn11* and *Mog*), and three proteins for astrocytes (*Slc6a9*, *Slc6a11* and *Kcnj10*). Due to the absence of highly correlated markers identified in original spatial proteomics data for microglia, they were excluded from the comparative analysis. The signature score for each cell type was calculated as log1p-transformed and averaged protein expressions, which subsequent used as the ground truth for cell-type composition to calculate the PCC and SPCC metrics with the predicted cell-type composition.

For the self-collected mouse PDAC data, the mIHC staining intensities from two image channels (*EpCAM* and *CD45*) were individually normalized to a range of 0 to 1, corresponding to PC and PI, respectively. This normalized intensity served as the ground truth and was utilized to calculate the PCC and SPCC metrics in comparison with the predicted cell-type composition of PC and PI.

To assess the reconstruction performance of Spatial-DC, we initially calculated the four metrics (CCC, RMSE, PCC and SPCC) between the synthetic and reconstructed spatially and cell-type resolved proteomic profiles. This involved several preprocessing steps: we filtered both the synthetic and reconstructed data to retain only cells with at least ten identified proteins and proteins identified in at least three cells. Following this, we used the log1p-normalized proteomic profiles of each cell type to compute the four metrics. To evaluate the capacity of identifying differential expression proteins (DEPs) using reconstructed data, we performed Welch’s two-sided t-tests on both the projection data and the reference data. Following this, we applied the Benjamini-Hochberg (BH) correction method to adjust the p-values for multiple comparisons. Proteins with an adjusted p-value less than 0.05 and a log2 fold change greater than 0 from both datasets were considered as DEPs. For visualization, we utilized the log1p-normalized and averaged abundance of DEPs from both the projection and reconstructed data to plot the heat map.

### Benchmark methods

To evaluate the deconvolution ability for estimating cell-type composition of spatial proteomics, we selected eight start-of-the-art methods for comparison, including Tangram, cell2location, Stereoscope, DestVI, Seurat, SpatialDWLS, SPOTlight and CARD. The running times of these methods are listed in **Table S9**.

Tangram (version 1.0.4) was employed following the tutorial (https://tangram-sc.readthedocs.io/en/latest/tutorial_sq_link.html). We used log1p to normalize the abundance of proteomics. Then we selected total 2,000 marker proteins for training. We use ‘map_cells_to_spacè function with ‘mode = clusters’ to find the optimal spatial alignment between reference proteomics data and spatial proteomics data.

Cell2location (version 0.1.3) was employed following the tutorial (https://cell2location.readthedocs.io/en/latest/notebooks/cell2location_tutorial.html). We trained a regression model to estimate the reference cell type signatures using original reference single-cell or single-cell-type proteomics data. Then we obtained the estimated expression of every protein for each cell type from the trained posterior distribution. Next, we created and trained the cell2location spatial mapping model with the default parameters, exporting 5% quantile estimated cell abundance. We trained 10,000 and 5,000 epochs for the reference data and target spatial proteomics data, respectively. The other parameters were set as same as in the tutorial.

Stereoscope (implemented in scvi-tools version 0.20.3) was employed following the tutorial (https://docs.scvi-tools.org/en/stable/tutorials/notebooks/spatial/stereoscope_heart_LV_tutorial.html). We trained a reference single-cell or single-cell-type model for 2,000 epochs, ensuring that the model was adequately optimized. Then, for each spatial dataset, we conducted a separate training of the spatial model for 2,000 epochs. The other parameters were used as default.

DestVI (implemented in scvi-tools version 0.20.3) was employed following the tutorial (https://docs.scvi-tools.org/en/stable/tutorials/notebooks/spatial/DestVI_tutorial.html). Similar to Stereoscope, we trained a reference single-cell or single-cell-type model for 2,000 epochs to ensure thorough optimization. Then, we conducted a separate training of the spatial model for 2,000 epochs. The other parameters were used as default.

Seurat (version 4.3.0.1) was employed following the tutorial (https://satijalab.org/seurat/articles/spatial_vignette). For data preprocessing, we performed ‘NormalizeDatà, ‘FindVariableFeatures’ and ‘ScaleDatà function sequentially with default parameters for both the reference and target spatial data, which served as the reference and query, respectively. For the deconvolution of each spot, we identified shared anchors using ‘FindTransferAnchors’ for the reference and query data with ‘k.score=10’, ‘npcs=5’ and ‘dims=1:5’ followed by transferring cell-type labels to query anchors using ‘TransferDatà function with ‘k.weight=20’. The predicted score of cell-type labels were converted to cell-type composition of each spot.

SPOTlight (version 1.2.0) was employed following the tutorial (https://marcelosua.github.io/SPOTlight/articles/SPOTlight_kidney.html). The preprocessing steps for both the reference and target spatial data followed the protocols akin to those employed in the Seurat. Specifically, the standardization and normalization processes were consistent with the practices in Seurat. To identify marker proteins within the reference data, the ‘FindAllMarkers’ function was utilized, applying its default parameters to comprehensively screen for distinguishing features.

SpatialDWLS (implemented in Giotto version 1.1.2) was employed following the tutorial (https://github.com/rdong08/spatialDWLS_dataset/blob/main/codes/Heart_ST_tutorial_spatialDWLS.Rmd). For the construction of signature expression matrix using reference cell-type proteomics data, up to top 1,000 ranked proteins were chosen. We created a Giotto object using the target spatial proteomics data without any filtering steps, followed by running the ‘runDWLSDeconv’ function for deconvolution each spot with default parameters. The deconvolution results stored in the Giotto object were then converted to cell-type composition.

CARD (version 1.0.0) was employed following the tutorial (https://yma-lab.github.io/CARD/documentation/04_CARD_Example.html). We created the CARD object using the original reference cell-type proteomics data and target spatial-proteomics data without any filtering, followed by performing deconvolution of each spot with default parameters.

### Generation of MS-based mouse PDAC spatial proteomics dataset

#### Animal experiments

In this study, we collected LMD-MS based spatial proteomics datasets from Kras^LSL-G12D/+^; Trp53^flox^; Pdx1-Cre mice (termed as KP^f/f^C mice), the mice were maintained at the appropriate biosafety level under constant temperature and humidity conditions in the Animal Experiments Center at Southern University of Science and Technology. Mouse organs were prepared as described previously^56,64^.

Nine-week-old KP^f/f^C mice were killed by cervical dislocation, and KP^f/f^C tumors of KP^f/f^C mice were collected and washed with ice-cold phosphate buffered saline (PBS) for three times. KP^f/f^C tumors were immersed in a 4% paraformaldehyde (PFA) solution for 24-48-hour fixation, and then, the tissues were paraffin-embedded for further processing. The animal experiments were approved by the Institutional Animal Care and Use Committee at Southern University of Science and Technology of China.

#### Multiplexed immunohistochemical (mIHC) staining

The formalin fixation and paraffin embedding (FFPE) KP^f/f^C tumors were sliced into sections with thickness of 4 μm by Leica RM 2255 microtome and mounted onto the MMI frame slides (MMI, 50103). For stronger tissue adhesion, the slides were dried at 55 °C for 30 minutes. Next, the slides were heated at 65 °C for 60 minutes, and were deparaffinized by washing 10 minutes in 100% xylene (Alfa Aescar) for three times, and followed by ethanol series (100%, 100%, 90%, 80%, and 70% (*v/v*) ethanol in water, each step for 5 min), and rehydrated by immersing in ddH_2_O for 5 minutes. TSA kit (TissueGnostics, TGFP7100) were used for the mIHC staining as the manufacturer’s instructions. Primary antibodies, anti-EpCAM (Cell Signaling Technology, clone E6V8Y, dilution 1:500) and anti-CD45 (Cell Signaling Technology, clone D3F8Q, dilution 1:1000) were used. Then, slides were washed three times for 2 minutes in PBS in the dark, and mounted with DAPI Fluoromount-G® antifade mountant (SouthernBiotech, 0100-20) and coverslips. Slides were imaged on TissueFAXS Spectra Systems (TissueGnostics) at 40× magnification.

#### Laser microdissection (LMD)

LMD was performed using a CellCut system (MMI). The cutting regions were defined by the rectangle tool in the MMI software, and were set as 100 μm×100 μm with an edge-to-edge distance of 10 μm. The following laser settings were used: 20x objective, brightfield mode, 10 μm/sec cut velocity, 165 μm laser focus, and 37.6% laser power. The cut regions were collected on the IsolationCap (MMI) and stored at −20 °C for further processing.

#### Sample preparation by iPAC

For protein extraction, 30 μL of DDM lysis buffer (1% (*w/v*) DDM, 10 mM HEPES (pH 7.4), 150 mM NaCl, 600 mM guanidine HCl, and protease inhibitor mixture (Roche)) was added to the IsolationCap (MMI) with tissue sections. The tissue sections were sonicated using Bioruptor (Diagenode) for 20 cycles (30 s-on, 30 s-off) at 4 °C, followed by heating at 95 °C for 90 minutes, and another round of sonication for 20 cycles. The iPAC spintip was assembled with 2 plugs of C18 disks (3M Empore) and 2 plugs of SAX disks (3M Empore) in 200 μL pipette tip. The iPAC spintips were activated with methanol and equilibrated with 3M ammonium hydroxide (NH_4_OH, pH 12) through centrifugation. Next, 30 μL of coating buffer (0.1% DDM in 6M NH_4_OH) was loaded into iPAC spintips by centrifugation at 50g for 5 seconds. Then, 30 μL of tissue lysates were mixed into the iPAC spintips with slow centrifugation for sufficient sample loading. The sample were washed with 20% (*v/v*) acetonitrile (ACN) in 3M NH_4_OH, followed by aggregated and captured at the SAX disks by incubating in pure ACN for 10 minutes, then further washing by 80% (*v/v*) ACN twice. After that, the samples were reduced by 50 mM dithiothreitol (DTT) in 20 mM ammonium bicarbonate (ABC) and incubated for 30 minutes at 37 °C. Followed by 20 mM ABC washing, the samples were digested using 4 μL of digestion buffer (10 mM iodoacetamide (IAA), 20 ng/μL sequencing-grade trypsin (Promega) and 20 ng/μL sequencing-grade Lys-C (Wako) in 20 mM ABC (pH 8.0)) and incubated for 3 hours at 37 °C in darkness. Next, 1 M NaCl in 1% formic acid (FA) was added to transfer the digested peptides to C18 disks, and 1% FA was loaded twice for desalting. The peptides were released from the C18 layer by washing with 80% ACN with 0.5% acetate into glass insert. The eluted peptides were lyophilized to dryness and dissolved in 0.1% (*v/v*) FA for MS analysis, 2/3 samples were injected.

#### Sample High-pH reversed-phase chromatography fractionation

High-pH reversed-phase chromatography fractionation was used to establish a deep proteomic library for data-independent analysis (DIA). The library of KP^f/f^C mouse tissues was generated in previous report^56^. Briefly, the whole mouse tumor was lysed, precipitated, and digested. One hundred micrograms of peptides from mouse brain were fractionated on a microflow HPLC (1260, Agilent) equipped with an XBridge peptide BEH C18 column (130 Å, 5 μm, 2.1 mm × 150 mm) within a 60 minute-gradient and concatenated into 24 fractions. All peptide samples were vacuum dried in a SpeedVac (Thermo Fischer), and re-dissolved in 0.1% FA spiked with 0.1X iRT peptides (Biognosys) for LC-MS/MS analysis.

#### LC-MS/MS analysis

All LC-MS/MS analysis were performed with a nanoElute UPLC system (Bruker Daltonik) connected to a trapped ion mobility spectrometry quadruple time-of-flight mass spectrometer (timsTOF Pro, Bruker Daltonik) via a captive spray ion source. The homemade 50 μm i.d. × 20 cm capillary column packed with C18 beads (1.9 μm, 120 Å, Dr. Maisch GmbH) was used for peptide separation. The column temperature was kept at 50 °C in an integrated column oven. Mobile phase A consists of 0.1% FA in water and mobile phase B consists of 0.1% FA in ACN. Peptides were separated using a linear gradient from 2 to 22% (*v/v*) mobile phase B in 50 minutes and stepped up to 35% (*v/v*) mobile phase B in 10 minutes, followed by an increase to 80% (*v/v*) mobile phase B in 10 minutes, and 10 minutes washing in 80% (*v/v*) mobile phase B at a flow rate of 100 nL/min.

For library establishment, ddaPASEF mode was adopted and the parameters were set as follow. The m/z scan range was set to 300-1500 in positive electrospray mode, and the 1/K0 scan range was set to 0.75-1.30 Vs/cm^2^. The ramp time was 200.0 ms and the total cycle time was 1.03 s with one MS1 scan and four PASEF MS/MS scans per acquisition cycle.

For data independent acquisition (DIA), the diaPASEF method was used as previous optimization^56^. Variable isolation windows adjusted by the precursor densities with m/z range 300-1500 and mobility range (1/K0) 0.75-1.30 Vs/cm^2^. The ramp time was 200.0 ms. The optimized method includes 2 ion mobility windows and 12 dia-PASEF scans. Total resulting DIA cycle time was estimated to be 2.71 s.

#### MS data analysis

For self-collected mouse PDAC spatial proteomics data, all MS raw files were searched against the reviewed mouse UniProt FASTA database (UP000005640, 21,984 entries) by the Spectronaut software (version 18.2, Biognosys) with all parameters set as default. The resulting library comprised 210,597 precursors accounting for 168,887 peptides and 11,438 protein groups. Next, a default library-based analysis pipeline was applied for protein identification and quantification of all DIA raw files.

### Spatial domain recognition

The K-means algorithm, sourced from the ‘sklearn.cluster’ module of the Python scikit-learn library (version 1.2.2), was employed to identify spatial domain. The input data comprised of spot-level proteomic profiles extracted from the spatial data and cell-type composition derived from deconvolution. Specifically, for the human palatine tonsil dataset, we normalized proteomic profiles using the log1p method and, alongside the cell-type composition for each spot, fed these into the K-means function, setting the number of clusters (n_clusters) to 3. For the PDAC data, we adjusted the n_clusters parameter to 2, while maintaining the other parameters at their default settings.

### Co-localization patterns analysis

This analysis began by leveraging the cell-type composition data derived from deconvolution methods for each spot. The cell-type composition data was used to calculate Pearson correlation coefficients, serving as a measure of the efficiency of co-localization between different cell types. Following this, Euclidean distance was computed based on the calculated co-localization efficiency, which was used to quantify the dissimilarity among the deconvoluted cell types.

### Mouse ligand-receptor database construction

To construct a cell-cell interaction network with highly confident PPIs, we created our mouse L-R interactions datasets with the following strategy. First, we created a database of mouse secreted and plasma proteins by downloading 1,556 protein entries with “GO-CC”, annotated with “extracellular space”, and 4,515 protein entries with “GO-CC”, annotated with “cell surface” or “plasma membrane” from UniProt database. Then, we downloaded two available mouse PPIs datasets, CellTalkDB^65^ and CellChatDB^66^. We retained only the protein pairs annotated with “secreted” in ligand protein and “plasma membrane” in receptor protein. This resulted in 742 and 825 protein pairs obtained from CellTalkDB and CellChatDB, respectively. Next, we downloaded CellPhoneDB_V5^67^ and obtained 1,218 homologous proteins. After integration of these interacted pairs, we constructed the database of mouse ligands and receptors, which contained 1,928 unique L-R interactions, comprising 554 ligand proteins and 457 receptor proteins. Subsequently, the curated database was utilized for L-R interactions identification as well as cell-cell interaction and communication analysis.

### Cell-cell communication analysis

For deciphering signals between sender and receiver cells from reconstructed spatially and cell-type resolved proteomic profiles, we employed the COMMOT^51^ algorithm with default parameters. For the human tonsil data and the PDAC data, we focused on inferring the paracrine cell-cell communications, which depend on the interaction of ligand proteins secreted from sender cells and receptor proteins expressed on the membrane of receiver cells. Consequently, we retained only ligand proteins from sender cells and receptor proteins from receiver cells, enabling us to screen signaling direction and strength. In contrast, for the mouse brain data, we identified the neuronal juxtacrine signaling, a mode of communication that necessitates direct cell-to-cell contact for the transfer of signaling molecules. The identified signaling proteins involved in these pathways were retained for both sender and receiver cells (**Table S6**).

### Survival time analysis of pancreatic patients

The survival analysis was carried out using the R package survival (version 3.5-5). Comprehensive proteomic abundance and associated clinical information of pancreatic patients were retrieved from LinkedOmicsKB^38^ (https://kb.linkedomics.org/download#PDAC). For the identification of the link between PPIs and the survival time of pancreatic patients, we first categorized patients into low-risk or high-risk groups based on the protein abundance of their ligand and receptor proteins. This categorization was achieved by employing the 50th percentile of protein abundance as the cutoff threshold. Specifically, patients with ligand and receptor protein abundances above this threshold were labeled as high-risk, whereas those below were labeled as low-risk. Subsequently, to ensure a focused analysis, we narrowed the PDAC patient cohort to include only individuals who were consistently classified as either high-risk or low-risk for both their ligand and receptor proteins. For the purpose of conducting a survival analysis centered on the *COL8A1-ITGA2* PPI, we selected a subgroup of 28 PDAC patients who exhibited relatively high abundance levels of this protein pair. Additionally, we identified a complementary group of 29 PDAC patients who displayed relatively low abundance levels of *COL8A1-ITGA2*. This approach allowed us to specifically investigate the potential impact of this interaction on PDAC patient outcomes within a well-defined and homogeneous patient population.

### Statistical Analysis

Statistical analysis of reconstructed cell-type proteomic profiles were performed using Perseus^68^ software (2.0.7.0) and R studio (4.1.0). MS-based proteomics data were imputed for deconvolution and statistical analysis, with missing values imputed by the strategy of one sample’s normal distribution of proteomic abundance using Perseus (down-shifted mean by 1.8 standard deviation and scaled by 0.3). The reconstructed cell-type samples with composition less than 0.1% from spatial proteomics were excluded for the following analysis. One-way ANOVA was performed for significance test of multiple sample and two-tailed Student’s t-test was used for calculating significance between two groups. clusterProfiler (3.16.1)^69^ in R was used for GO enrichment analysis.

## Data availability

All synthetic and real-world proteomics datasets analyzed in this study, including their source links or accession codes and pre-processed steps, are provided in **Note S1**. The details of synthetic datasets and real-world datasets are summarized in **Table S2** and **Table S3**, respectively. The Mouse PDAC dataset was generated specifically for this study. The MS raw files have been deposited in the ProteomeXchange Consortium via the PRIDE partner repository and can be accessed with the dataset identifier IPX0007607001 [https://www.iprox.cn/page/SSV024.html;url=17337921146112PmZ] (Password: w9pl). The processed reference and spatial proteomics datasets, along with the processed mIHC imaging data of mouse PDAC tissue are available on Zenodo [https://zenodo.org/records/14233847]. All other data supporting the findings of this study are included in the Supplementary Information.

## Code availability

The source code for pre-processing, data analysis, and generating figure and tables has been deposited in both GitHub [https://github.com/TencentAILabHealthcare/Spatial-DC] and Zenodo [https://doi.org/10.5281/zenodo.14386585].

## Acknowledgements

R.T. is approved by the funding from China State Key Basic Research Program Grants (2024YFA1307200, 2021YFA1301602, 2022YFC3401104, 2021YFA1301601, 2021YFA1302603), National Natural Science Foundation of China (92253304, 22125403, 32201218, 82341074, and 22104047), Shenzhen Innovation of Science and Technology Commission (JSGGZD20220822095200001, JCYJ20210324120210029, and KJZD20230923114220041), Shenzhen Science and Technology Program (ZDSYS20230626090803004).

## Author contributions

Y.L., Q.K. and Z.W. conceived the deconvolution for spatial proteomics concept, devised strategies for associated experiments, analyzed the data, and wrote the paper. Z.W. and Y.L. developed and implemented Spatial-DC. Q.K. and X.W. conducted mIHC imaging staining and LMD-MS based spatial proteomics experiments for mouse PDAC data. Y.X., Y.M. and X.W. aided in the collection and analysis of mouse PDAC tissue. Y.G. offered advice on adjusting the parameters of Spatial-DC, and W.G. provided analysis strategies for the spatial proteomics data. J.Y. and R.T. contributed to refining the concept of Spatial-DC, offered advice on experiment design and analysis strategies, and assisted in revising the paper. All authors engaged in discussions about the results and implications, as well as provided feedback on the manuscript throughout its development.

## References

1. Cao, L. et al. Proteogenomic characterization of pancreatic ductal adenocarcinoma. Cell 184, 5031–5052.e26 (2021).

2. Luca, B. A. et al. Atlas of clinically distinct cell states and ecosystems across human solid tumors. Cell 184, 5482–5496.e28 (2021).

3. Ribas, A. & Wolchok, J. D. Cancer immunotherapy using checkpoint blockade. Science (2018) doi:10.1126/science.aar4060.

4. Zhang, L. et al. Single-Cell Analyses Inform Mechanisms of Myeloid-Targeted Therapies in Colon Cancer. Cell 181, 442–459.e29 (2020).

5. Grünwald, B. T. et al. Spatially confined sub-tumor microenvironments in pancreatic cancer. Cell 184, 5577–5592.e18 (2021).

6. Galon, J. & Bruni, D. Approaches to treat immune hot, altered and cold tumours with combination immunotherapies. Nat Rev Drug Discov 18, 197–218 (2019).

7. Ståhl, P. L. et al. Visualization and analysis of gene expression in tissue sections by spatial transcriptomics. Science 353, 78–82 (2016).

8. Baysoy, A., Bai, Z., Satija, R. & Fan, R. The technological landscape and applications of single-cell multi-omics. Nat Rev Mol Cell Biol 24, 695–713 (2023).

9. Vandereyken, K., Sifrim, A., Thienpont, B. & Voet, T. Methods and applications for single-cell and spatial multi-omics. Nat Rev Genet 24, 494–515 (2023).

10. Rodriques, S. G. et al. Slide-seq: A scalable technology for measuring genome-wide expression at high spatial resolution. Science 363, 1463–1467 (2019).

11. Chen, A. et al. Single-cell spatial transcriptome reveals cell-type organization in the macaque cortex. Cell 186, 3726–3743.e24 (2023).

12. Mund, A., Brunner, A.-D. & Mann, M. Unbiased spatial proteomics with single-cell resolution in tissues. Molecular Cell 82, 2335–2349 (2022).

13. Liu, Y., Beyer, A. & Aebersold, R. On the Dependency of Cellular Protein Levels on mRNA Abundance. Cell 165, 535–550 (2016).

14. Ben-Chetrit, N. et al. Integration of whole transcriptome spatial profiling with protein markers. Nat Biotechnol 41, 788–793 (2023).

15. Method of the Year 2024: spatial proteomics. Nat Methods 21, 2195–2196 (2024).

16. Rieckmann, J. C. et al. Social network architecture of human immune cells unveiled by quantitative proteomics. Nat Immunol 18, 583–593 (2017).

17. Mund, A. et al. Deep Visual Proteomics defines single-cell identity and heterogeneity. Nat Biotechnol 40, 1231–1240 (2022).

18. Schürch, C. M. et al. Coordinated Cellular Neighborhoods Orchestrate Antitumoral Immunity at the Colorectal Cancer Invasive Front. Cell 182, 1341–1359.e19 (2020).

19. Mao, Y., Wang, X., Huang, P. & Tian, R. Spatial proteomics for understanding the tissue microenvironment. Analyst 146, 3777–3798 (2021).

20. Liu, Y. et al. High-plex protein and whole transcriptome co-mapping at cellular resolution with spatial CITE-seq. Nat Biotechnol 41, 1405–1409 (2023).

21. Ma, M. et al. In-depth mapping of protein localizations in whole tissue by micro-scaffold assisted spatial proteomics (MASP). Nat Commun 13, 7736 (2022).

22. Li, B. et al. Benchmarking spatial and single-cell transcriptomics integration methods for transcript distribution prediction and cell type deconvolution. Nat Methods 19, 662–670 (2022).

23. Li, H. et al. A comprehensive benchmarking with practical guidelines for cellular deconvolution of spatial transcriptomics. Nat Commun 14, 1548 (2023).

24. Garmire, L. X. et al. Challenges and perspectives in computational deconvolution of genomics data. Nat Methods (2024) doi:10.1038/s41592-023-02166-6.

25. Kleshchevnikov, V. et al. Cell2location maps fine-grained cell types in spatial transcriptomics. Nat Biotechnol 40, 661–671 (2022).

26. Andersson, A. et al. Single-cell and spatial transcriptomics enables probabilistic inference of cell type topography. Commun Biol 3, 565 (2020).

27. Lopez, R. et al. DestVI identifies continuums of cell types in spatial transcriptomics data. Nat Biotechnol 40, 1360– 1369 (2022).

28. Elosua-Bayes, M., Nieto, P., Mereu, E., Gut, I. & Heyn, H. SPOTlight: seeded NMF regression to deconvolute spatial transcriptomics spots with single-cell transcriptomes. Nucleic Acids Research 49, e50–e50 (2021).

29. Ma, Y. & Zhou, X. Spatially informed cell-type deconvolution for spatial transcriptomics. Nat Biotechnol 40, 1349– 1359 (2022).

30. Dong, R. & Yuan, G.-C. SpatialDWLS: accurate deconvolution of spatial transcriptomic data. Genome Biol 22, 145 (2021).

31. Biancalani, T. et al. Deep learning and alignment of spatially resolved single-cell transcriptomes with Tangram. Nat Methods 18, 1352–1362 (2021).

32. Stuart, T. et al. Comprehensive Integration of Single-Cell Data. Cell 177, 1888–1902.e21 (2019).

33. Vallania, F. et al. Leveraging heterogeneity across multiple datasets increases cell-mixture deconvolution accuracy and reduces biological and technical biases. Nat Commun 9, 4735 (2018).

34. Deep learning–based cell composition analysis from tissue expression profiles. SCIENCE ADVANCES (2020).

35. Chen, Y. et al. Deep autoencoder for interpretable tissue-adaptive deconvolution and cell-type-specific gene analysis. Nat Commun 13, 6735 (2022).

36. Xun, Z. et al. Reconstruction of the tumor spatial microenvironment along the malignant-boundary-nonmalignant axis. Nat Commun 14, 933 (2023).

37. Newman, A. M. et al. Determining cell type abundance and expression from bulk tissues with digital cytometry. Nat Biotechnol 37, 773–782 (2019).

38. Y, L. et al. A proteogenomics data-driven knowledge base of human cancer. Cell systems 14, (2023).

39. Song, D. et al. scDesign3 generates realistic in silico data for multimodal single-cell and spatial omics. Nat Biotechnol 42, 247–252 (2024).

40. Sun, T., Song, D., Li, W. V. & Li, J. J. scDesign2: a transparent simulator that generates high-fidelity single-cell gene expression count data with gene correlations captured. Genome Biol 22, 163 (2021).

41. Zhu, J., Shang, L. & Zhou, X. SRTsim: spatial pattern preserving simulations for spatially resolved transcriptomics. Genome Biol 24, 39 (2023).

42. Dou, M. et al. High-Throughput Single Cell Proteomics Enabled by Multiplex Isobaric Labeling in a Nanodroplet Sample Preparation Platform. Anal. Chem. 91, 13119–13127 (2019).

43. Woo, J. et al. High-throughput and high-efficiency sample preparation for single-cell proteomics using a nested nanowell chip. Nat Commun 12, 6246 (2021).

44. Massoni-Badosa, R. et al. An atlas of cells in the human tonsil. Immunity 57, 379–399.e18 (2024).

45. Makhmut, A. et al. A framework for ultra-low-input spatial tissue proteomics. Cell Systems 14, 1002–1014.e5 (2023).

46. De Silva, N. S. & Klein, U. Dynamics of B cells in germinal centres. Nat Rev Immunol 15, 137–148 (2015).

47. Nave, H., Gebert, A. & Pabst, R. Morphology and immunology of the human palatine tonsil. Anatomy and Embryology 204, 367–373 (2001).

48. Jhaveri, N. et al. Mapping the Spatial Proteome of Head and Neck Tumors: Key Immune Mediators and Metabolic Determinants in the Tumor Microenvironment. GEN Biotechnology 2, 418–434 (2023).

49. De Silva, N. S. & Klein, U. Dynamics of B cells in germinal centres. Nat Rev Immunol 15, 137–148 (2015).

50. Heesters, B. A., Myers, R. C. & Carroll, M. C. Follicular dendritic cells: dynamic antigen libraries. Nat Rev Immunol 14, 495–504 (2014).

51. Cang, Z. et al. Screening cell–cell communication in spatial transcriptomics via collective optimal transport. Nat Methods 20, 218–228 (2023).

52. Sharma, K. et al. Cell type– and brain region–resolved mouse brain proteome. Nat Neurosci 18, 1819–1831 (2015).

53. Crossin, K. L. & Krushel, L. A. Cellular signaling by neural cell adhesion molecules of the immunoglobulin superfamily. Dev. Dyn. 218, 260–279 (2000).

54. Ditlevsen, D. K., Povlsen, G. K., Berezin, V. & Bock, E. NCAM-induced intracellular signaling revisited. J of Neuroscience Research 86, 727–743 (2008).

55. Damo, E. & Simonetti, M. Axon Guidance Molecules and Pain. (2022) doi:10.3390/cells11193143.

56. Xu, Y. et al. Multimodal single cell-resolved spatial proteomics reveals pancreatic tumor heterogeneity. 2023.11.04.565590 Preprint at 10.1101/2023.11.04.565590 (2023).

57. Huang, P., Gao, W., Fu, C. & Tian, R. Functional and Clinical Proteomic Exploration of Pancreatic Cancer. Molecular & Cellular Proteomics 22, 100575 (2023).

58. Hou, G., Mulholland, D., Gronska, M. A. & Bendeck, M. P. Type VIII Collagen Stimulates Smooth Muscle Cell Migration and Matrix Metalloproteinase Synthesis after Arterial Injury. The American Journal of Pathology 156, 467– 476 (2000).

59. Yang, K. D. et al. Multi-domain translation between single-cell imaging and sequencing data using autoencoders. Nat Commun 12, 31 (2021).

60. Yang, F. et al. scBERT as a large-scale pretrained deep language model for cell type annotation of single-cell RNA-seq data. Nat Mach Intell 4, 852–866 (2022).

61. Cui, H. et al. scGPT: toward building a foundation model for single-cell multi-omics using generative AI. Nat Methods (2024) doi:10.1038/s41592-024-02201-0.

62. Lazar, C., Gatto, L., Ferro, M., Bruley, C. & Burger, T. Accounting for the Multiple Natures of Missing Values in Label-Free Quantitative Proteomics Data Sets to Compare Imputation Strategies. J. Proteome Res. 15, 1116–1125 (2016).

63. Kipf, T. N. & Welling, M. Semi-Supervised Classification with Graph Convolutional Networks. Preprint at http://arxiv.org/abs/1609.02907 (2017).

64. Huang, P., Li, H., Gao, W., Cai, Z. & Tian, R. A Fully Integrated Spintip-Based Approach for Sensitive and Quantitative Profiling of Region-Resolved in Vivo Brain Glycoproteome. Analytical Chemistry (2019) doi:10.1021/acs.analchem.9b01930.

65. Shao, X. et al. CellTalkDB: a manually curated database of ligand–receptor interactions in humans and mice. Briefings in Bioinformatics 22, bbaa269 (2021).

66. Jin, S. et al. Inference and analysis of cell-cell communication using CellChat. Nat Commun 12, 1088 (2021).

67. Troulé, K. et al. CellPhoneDB v5: inferring cell-cell communication from single-cell multiomics data.

68. Tyanova, S. & Cox, J. Perseus: A Bioinformatics Platform for Integrative Analysis of Proteomics Data in Cancer Research. in Cancer Systems Biology (ed. Von Stechow, L.) vol. 1711 133–148 (Springer New York, New York, NY, 2018).

69. Yu, G., Wang, L.-G., Han, Y. & He, Q.-Y. clusterProfiler: an R Package for Comparing Biological Themes Among Gene Clusters. OMICS: A Journal of Integrative Biology 16, 284–287 (2012).

70. Ma, M. et al. In-depth mapping of protein localizations in whole tissue by micro-scaffold assisted spatial proteomics (MASP). Nat Commun 13, 7736 (2022).

