## Supplementary Information for "Spatial-DC: a robust deep learning-based method for deconvolution of spatial proteomics"

### Supplementary Figures

#### Fig. S1

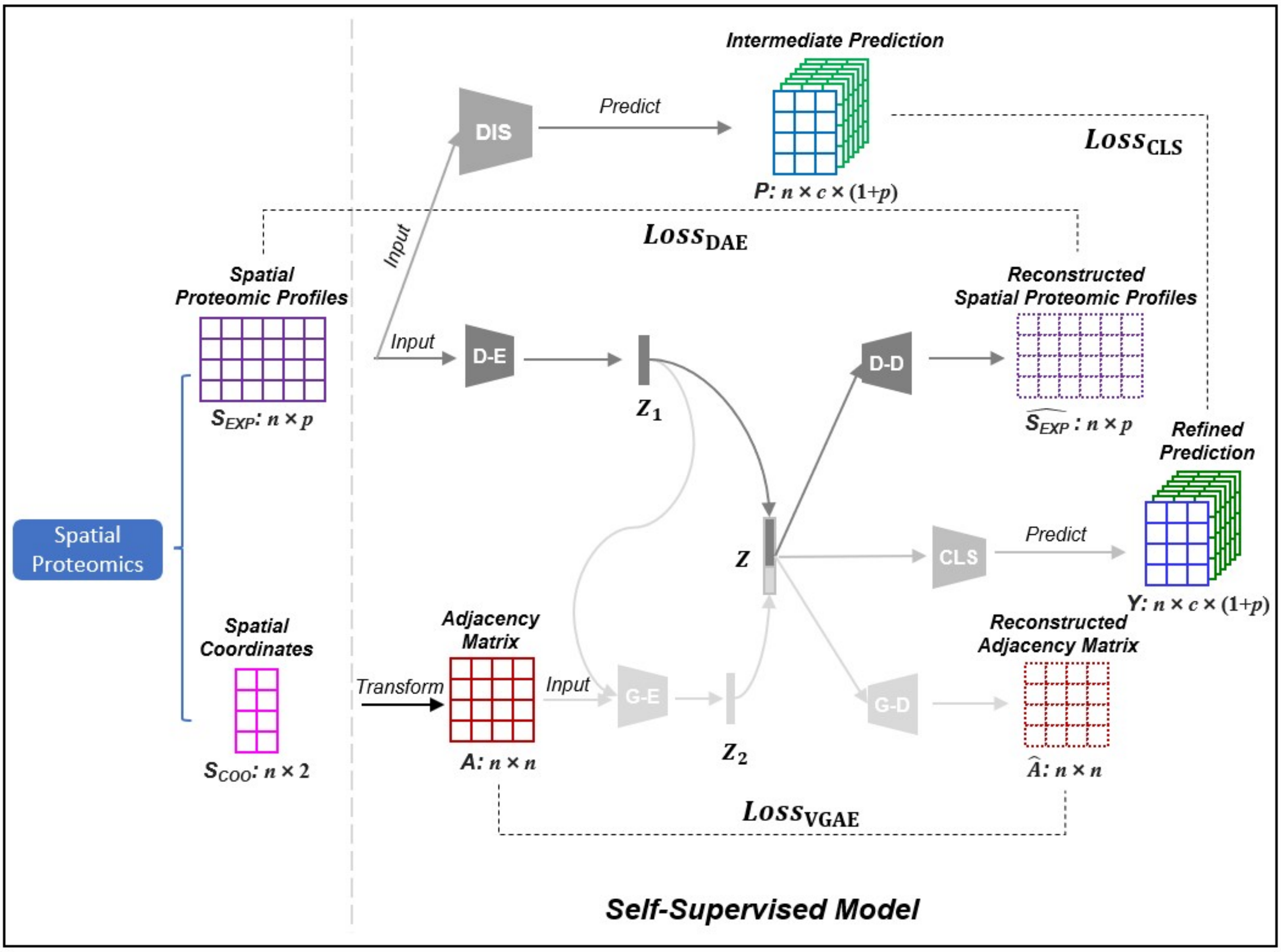


**Fig. S1 | Architecture of the self-supervised model in Spatial-DC.** The self-supervised model leverages the proteomic profiles and spatial coordinates of each spot to refine the intermediate prediction of cell-type resolved information, resulting in refined cell-type resolved information based on characteristics of the spatial proteomics data, which are subsequently converted to cell-type composition and cell-type resolved proteomic profiles for each spot (**Methods**). The self-supervised model comprises three main components: a deep autoencoder (**DAE**), a variational graph autoencoder (**VGAE**), and a classifier (**CLS**). **DAE**, comprising an encoder (**D-E**) and a decoder (**D-D**), is used to learn the compressed representation $Z_1$ of proteomic profiles $S_{EXP}$ of spatial proteomics data. **VGAE**, composed of a graph convolutional network (GCN) encoder (**G-E**) and an inner product decoder (**G-D**), is used to learn the latent graph-structured representations $Z_2$ of spatial proteomics data. **CLS** is a deep neural network (DNN) that processes the concatenated embedding $Z$ to generate the final prediction.

#### Fig. S2

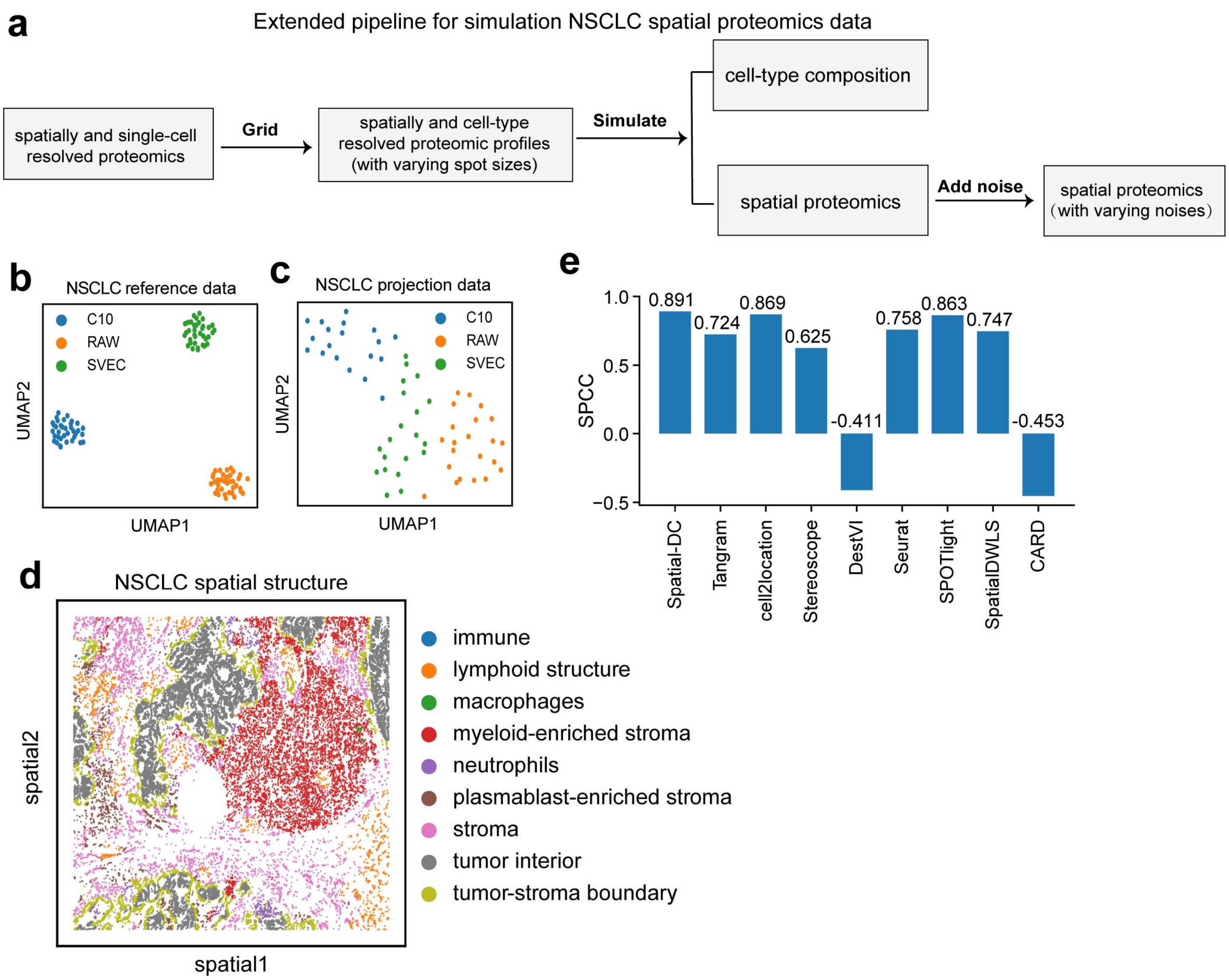


**Fig. S2 | Supplementary information of synthetic NSCLC spatial proteomics data. a,** The pipeline providing the external details for obtaining synthetic spatial proteomics data (**Methods**). The spatially and cell-type resolved proteomic profiles were used as the ground truth for reconstruction, while the cell-type composition served as the ground truth for deconvolution. **b-c**, UMAP plots of NSCLC reference data[1] (consisting of 1,437 proteins for 108 cells) and projection data[2] (consisting of 1,225 proteins for 61 cells). **d**, Spatial map demonstrating the spatial structure of NSCLC data, which records the relative spatial location of 32,859 cells across nine niches, each represented by a different color. **e,** Quantitative assessment by SPCC metrics of Spatial-DC performance on representative synthetic spatial proteomics data (with spot size of 100 μm). The results are presented by the mean values obtained from three cell types.

### Fig. S3

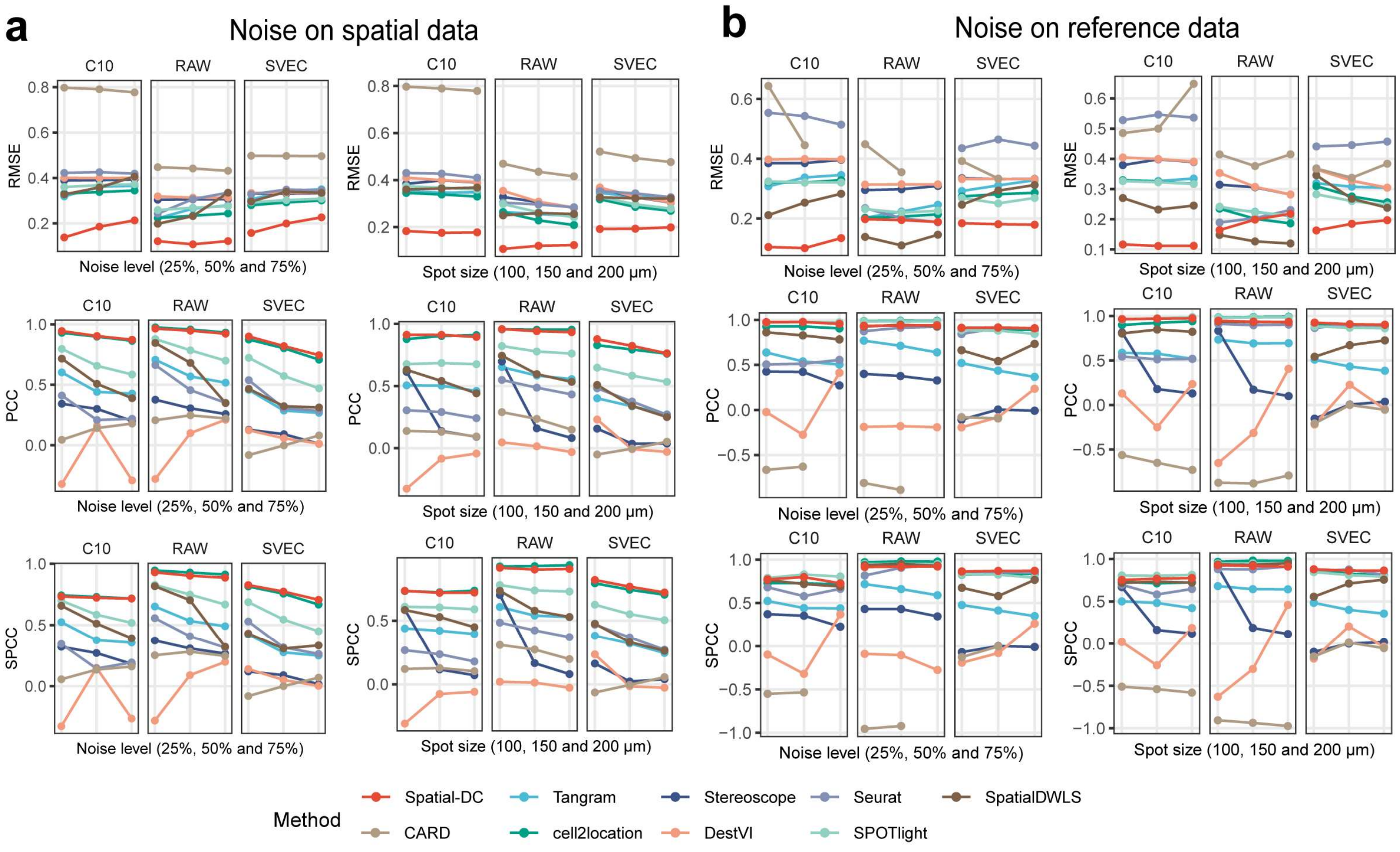


**Fig. S3 | Robustness evaluation of Spatial-DC for each cell type on synthetic NSCLC spatial proteomics data. a-b,** Evaluation metrics (RMSE, PCC and SPCC) were utilized to assess the robustness of Spatial-DC for each cell type on synthetic NSCLC data under varying levels of external noise on spatial data (**a**) and reference data (**b**). Line plots in **a** and **b** show the three metrics of Spatial-DC and eight state-of-the-art deconvolution methods across three cell types (left, middle and right) under varying noise levels (left column) and different spot sizes (right column). Each data point, corresponding to varying noise levels (left column), represents the mean metrics (RMSE, PCC and SPCC) from three spot sizes, and vice versa (**Table S4**).

### Fig. S4

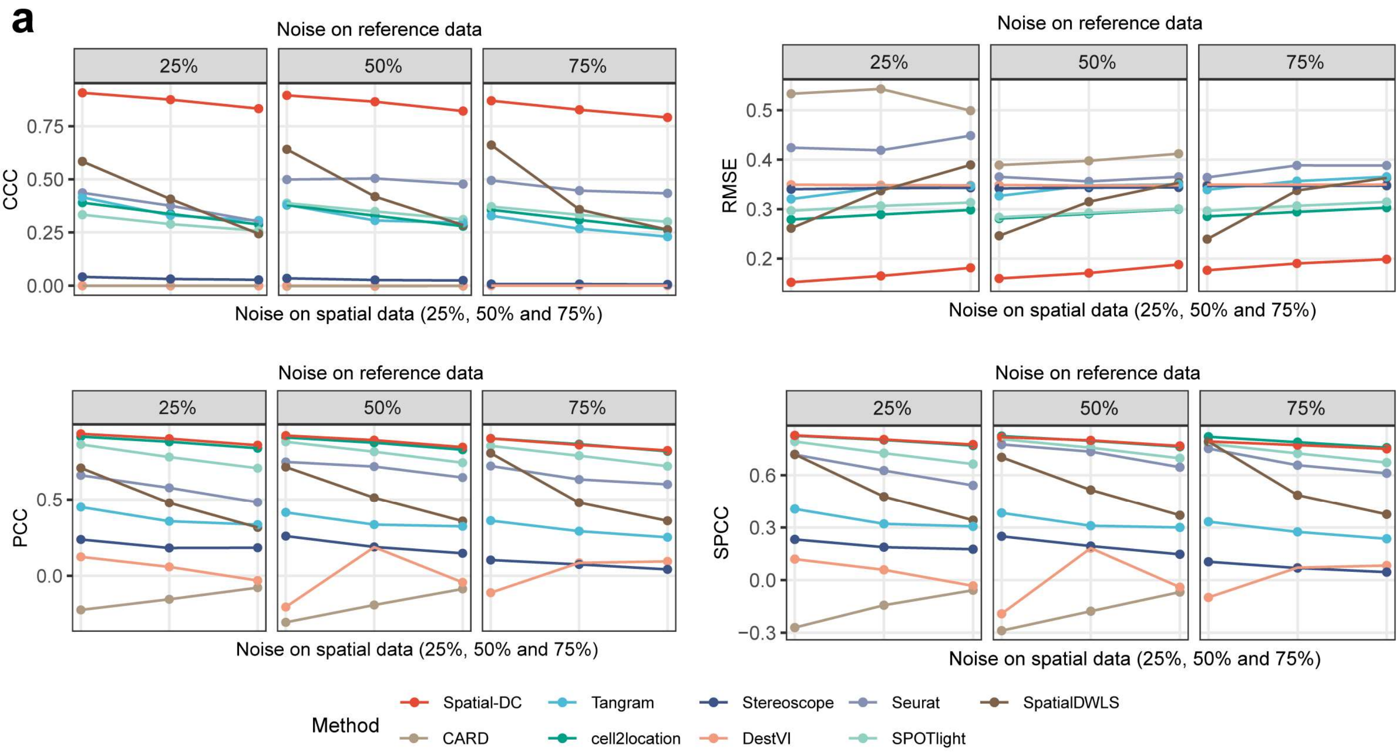


**Fig. S4 | Robustness evaluation of Spatial-DC for noise disturbance on synthetic NSCLC spatial proteomics data. a,** Evaluation metrics (CCC, RMSE, PCC and SPCC) were utilized to assess the robustness of Spatial-DC on synthetic NSCLC data, with external Gaussian noise added to both reference and spatial proteomics data. Each data point represents the mean metrics (RMSE, PCC and SPCC) derived from three cell types (RAW, SVEC and C10) under three different spot sizes (100, 150 and 200 μm) (**Table S4**).

### Fig. S5

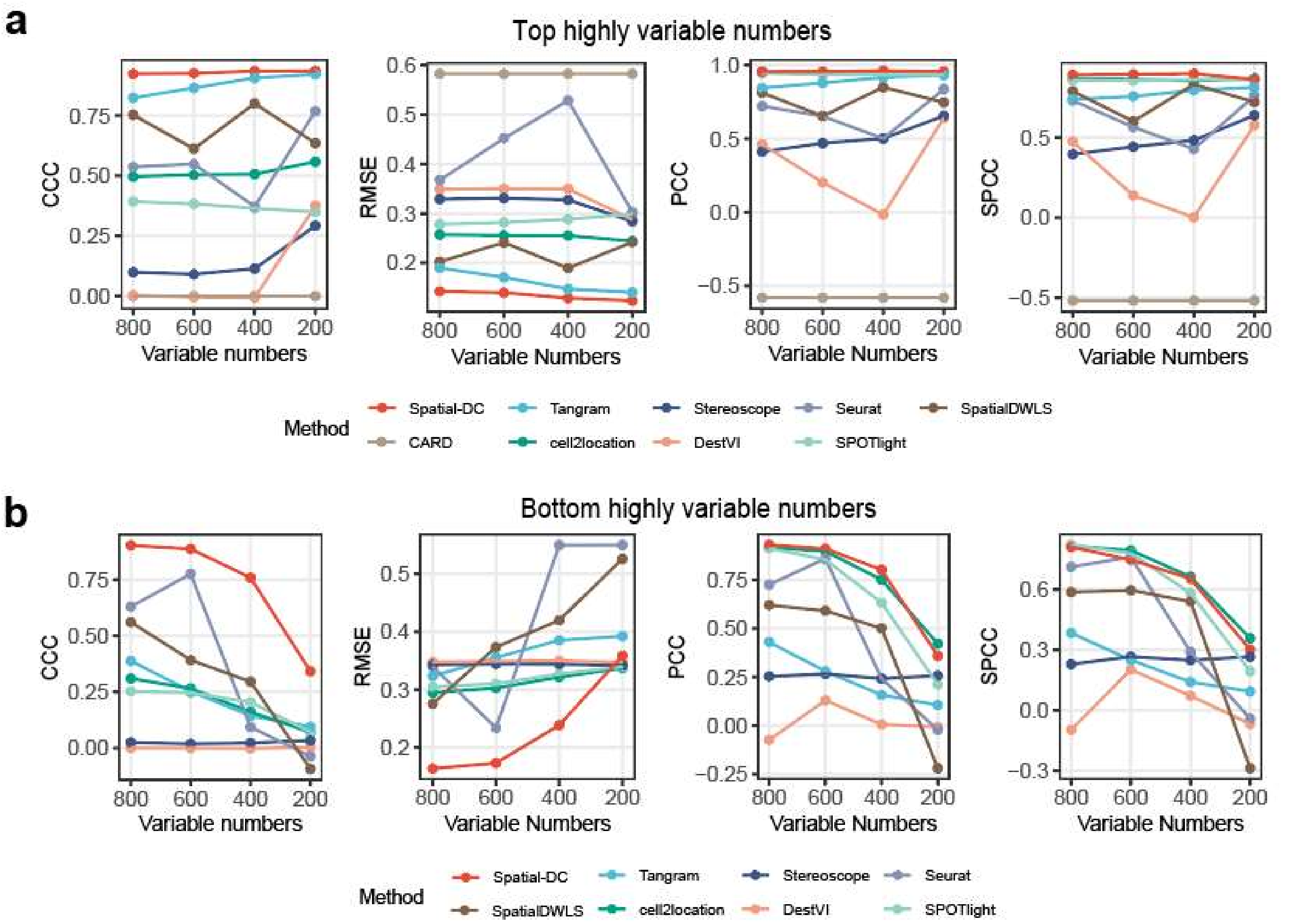


**Fig. S5 | Robustness evaluation of Spatial-DC for the protein panel used for deconvolution on synthetic NSCLC spatial proteomics data. a-b,** Evaluation metrics (CCC, RMSE, PCC and SPCC) were utilized to assess the robustness of Spatial-DC on synthetic NSCLC data, using the different protein panels that defined as top variable (**a**) and bottom variable (**b**) (**Methods**). Each data point represents the mean metrics (RMSE, PCC and SPCC) derived from three cell types (RAW, SVEC and C10) under three different spot sizes (100, 150 and 200 μm) (**Table S4**).

#### Fig. S6

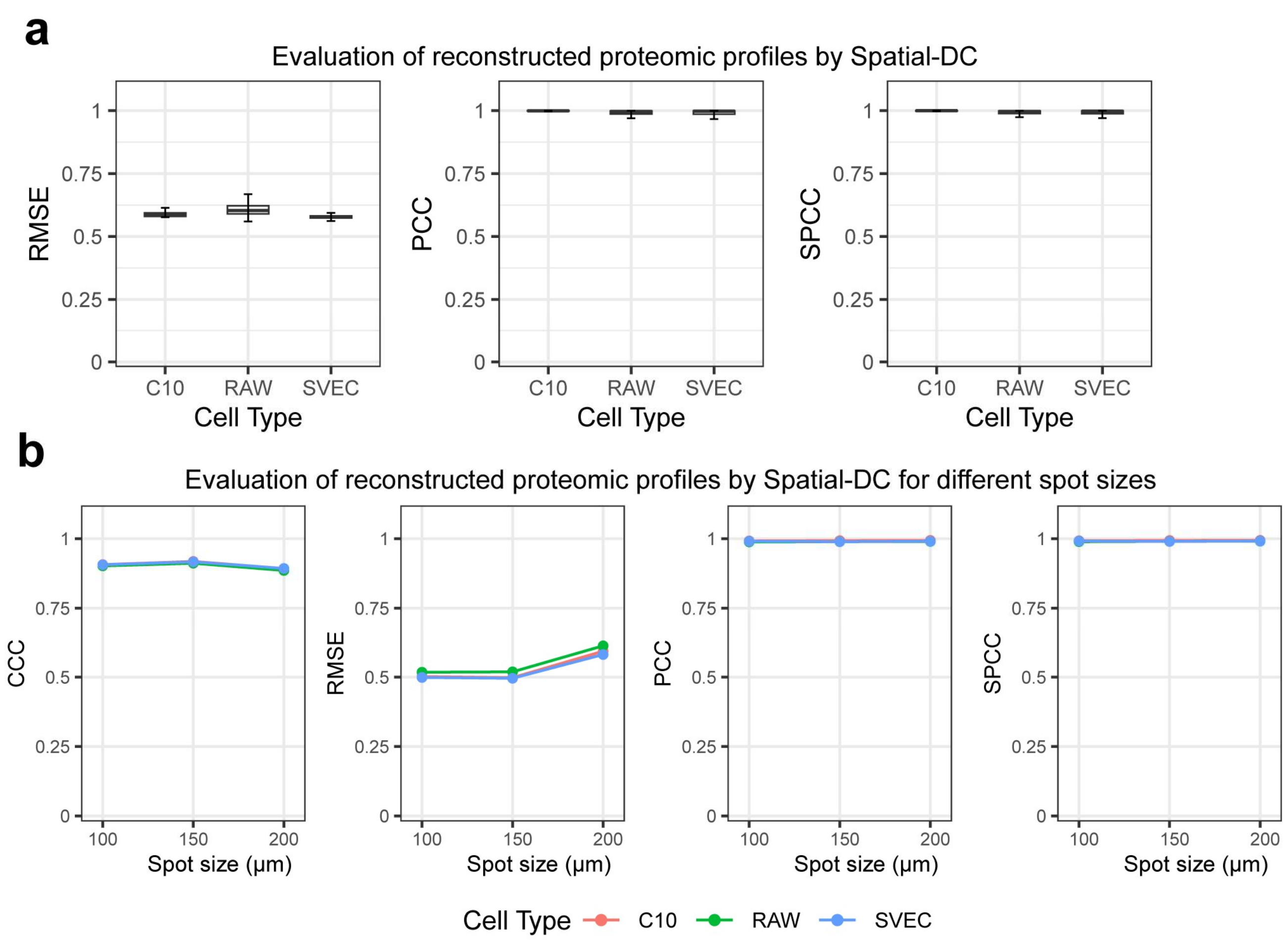


**Fig. S6 | Effectiveness evaluation of reconstructed cell-type resolved proteomic profiles by Spatial-DC on synthetic NSCLC spatial proteomics data. a,** Quantitative assessment by RMSE, PCC and SPCC metrics of reconstructed cell-type resolved proteomic profiles by Spatial-DC for three cell types from representative synthetic NSCLC data (with spot size of 200 μm). **b,** Quantitative assessment by CCC, RMSE, PCC and SPCC metrics of reconstructed cell-type resolved proteomic profiles by Spatial-DC for three cell types under varying sizes of synthetic spot.

#### Fig. S7

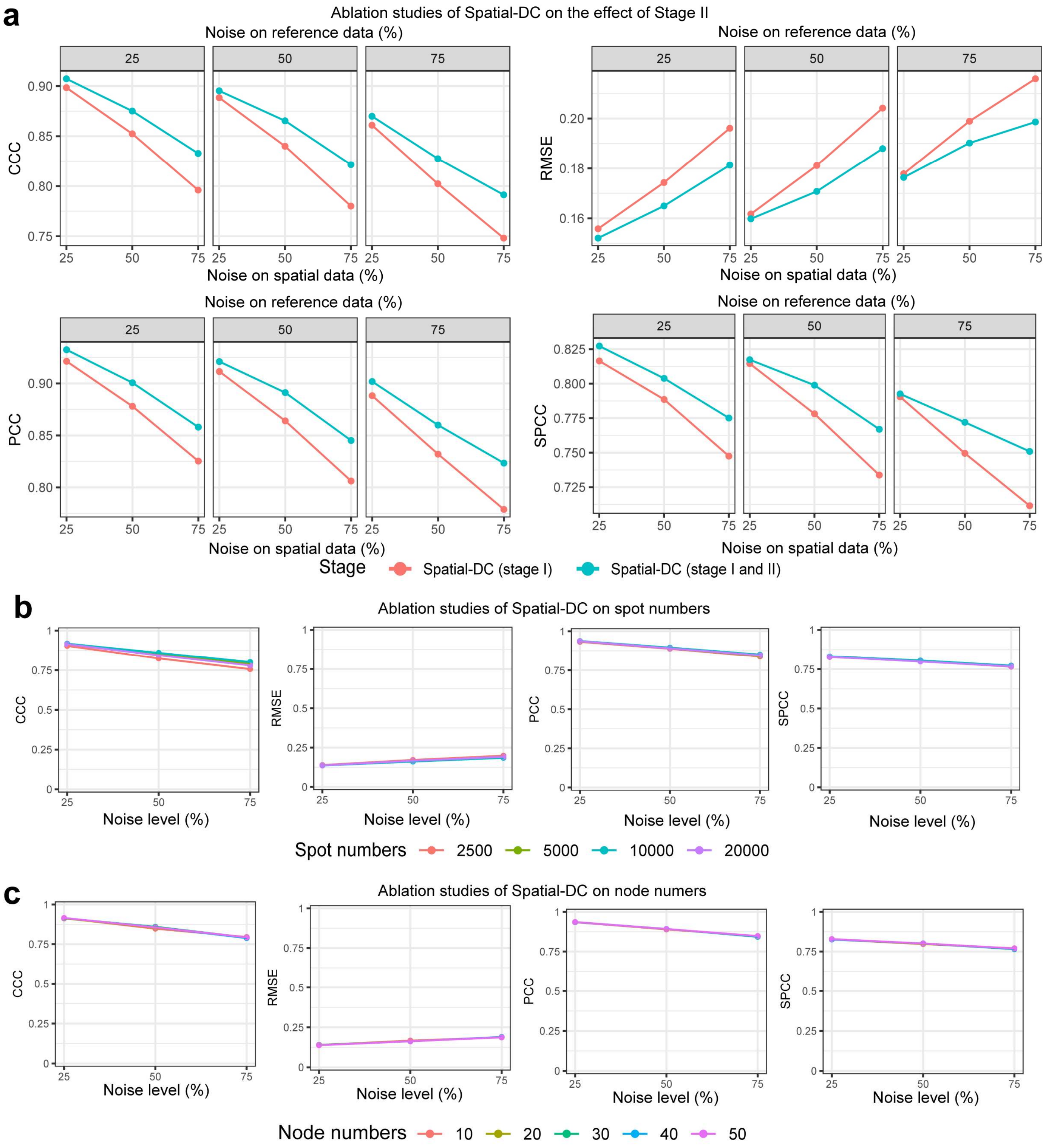


**Fig. S7 | Ablation studies of Spatial-DC on synthetic NSCLC spatial proteomics data. a-c,** Ablation studies of Spatial-DC under various noise levels with different parameters or components, including the inclusion of Stage II, the number of simulated spots used for training the distribution model **DIS**, and the number of nodes used to construct the nearest neighbor graph. **a,** Line plot comparing the CCC metrics with or without the inclusion of stage II of Spatial-DC on synthetic NSCLC spatial proteomics data. **b,** Line plot showing the CCC metrics achieved when using different numbers of simulated spots for training the **DIS** model. **c,** Line plot presenting the CCC values obtained when using different numbers of graph nodes to construct the nearest neighbor graph.

### Fig. S8

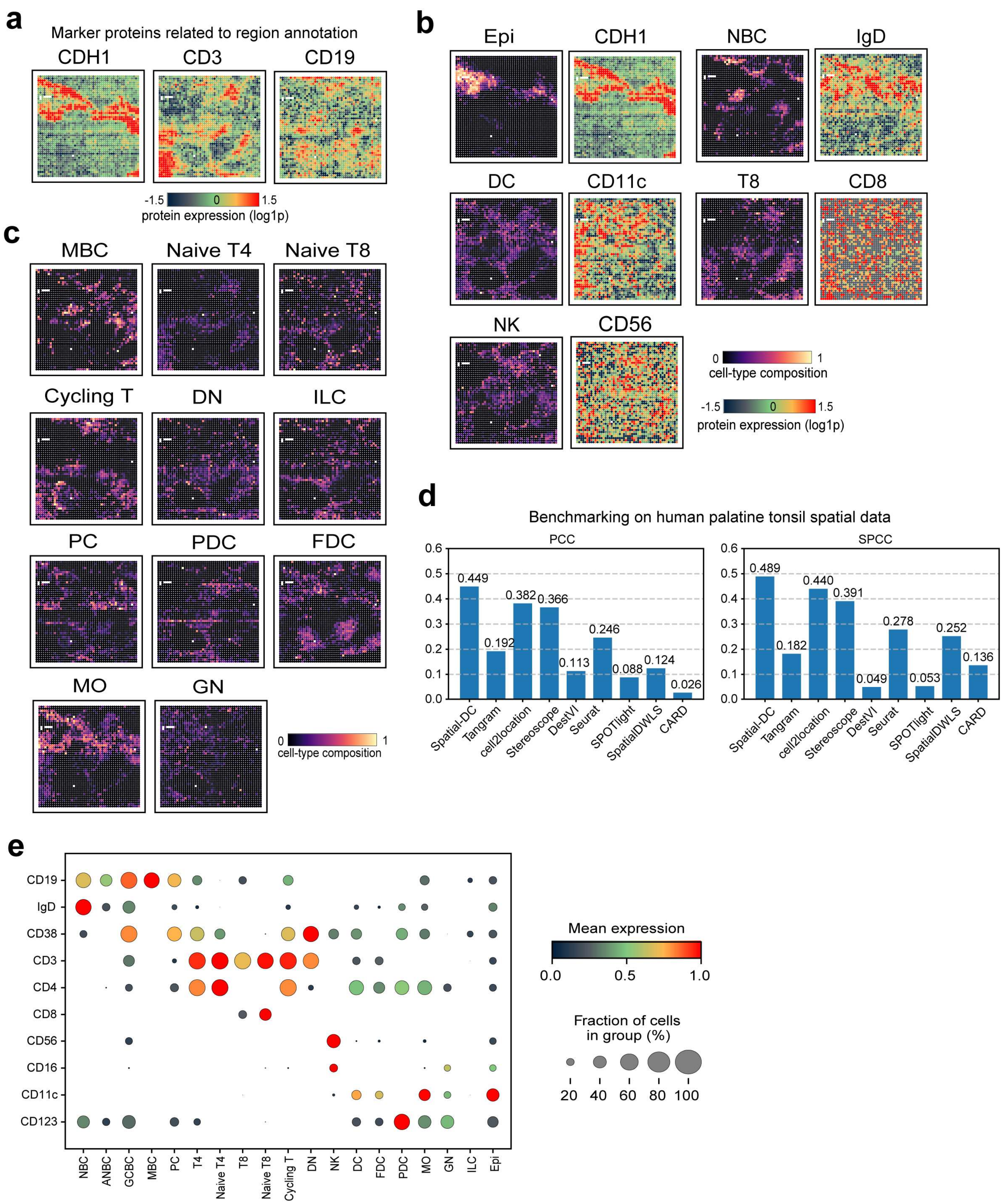


**Fig. S8 | Supplementary information on the application of Spatial-DC on human palatine tonsil data. a,** Spatial map of the known cell-type specific markers from human palatine tonsil spatial proteomics data. **b,**

Visualization of cell-type distributions predicted by Spatial-DC (left) and corresponding cell-type-specific markers identified from tonsil spatial proteomics data (right), as a supplement to **Fig. 3e**. **c,** Spatial map showing the rest cell-type distributions predicted from Spatial-DC without corresponding cell-type-specific markers, as a supplement to **Fig. 3e and b**. **d,** Bar plots depicting the mean PCC (left) and SPCC (right) metrics between deconvolution predictions and scaled abundance of markers of pan-B cells (*CD19*), pan-T cells (*CD3*), and epithelial cells (*CDH1*). **e,** Dot plot showing the mean expression of known cell-type-specific marker proteins from reconstructed cell-type proteomic profiles (**Table S10**), with the size of dot representing the fraction of cells in reconstructed spots.

#### Fig. S9

a

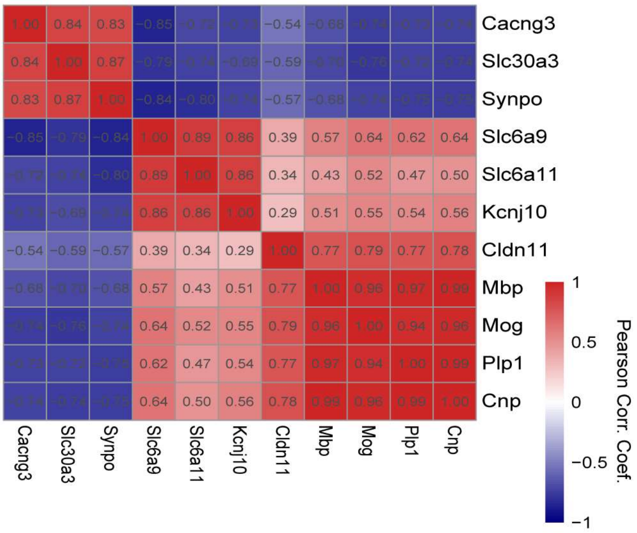


**Fig. S9 | Supplementary information on the application of Spatial-DC on mouse brain coronal data. a,** Heat map showing the Pearson correlation coefficient between cell-type-specific markers (**Methods**), including *Cldn11*, *Mbp*, *Mog*, *Plp1* and *Cnp* for oligodendrocytes, *Slc6a9*, *Slc6a11*, *Kcnj10* for astrocytes and *Cacng3*, *Slc30a3*, *Synpo* for neurons.

#### Fig. S10

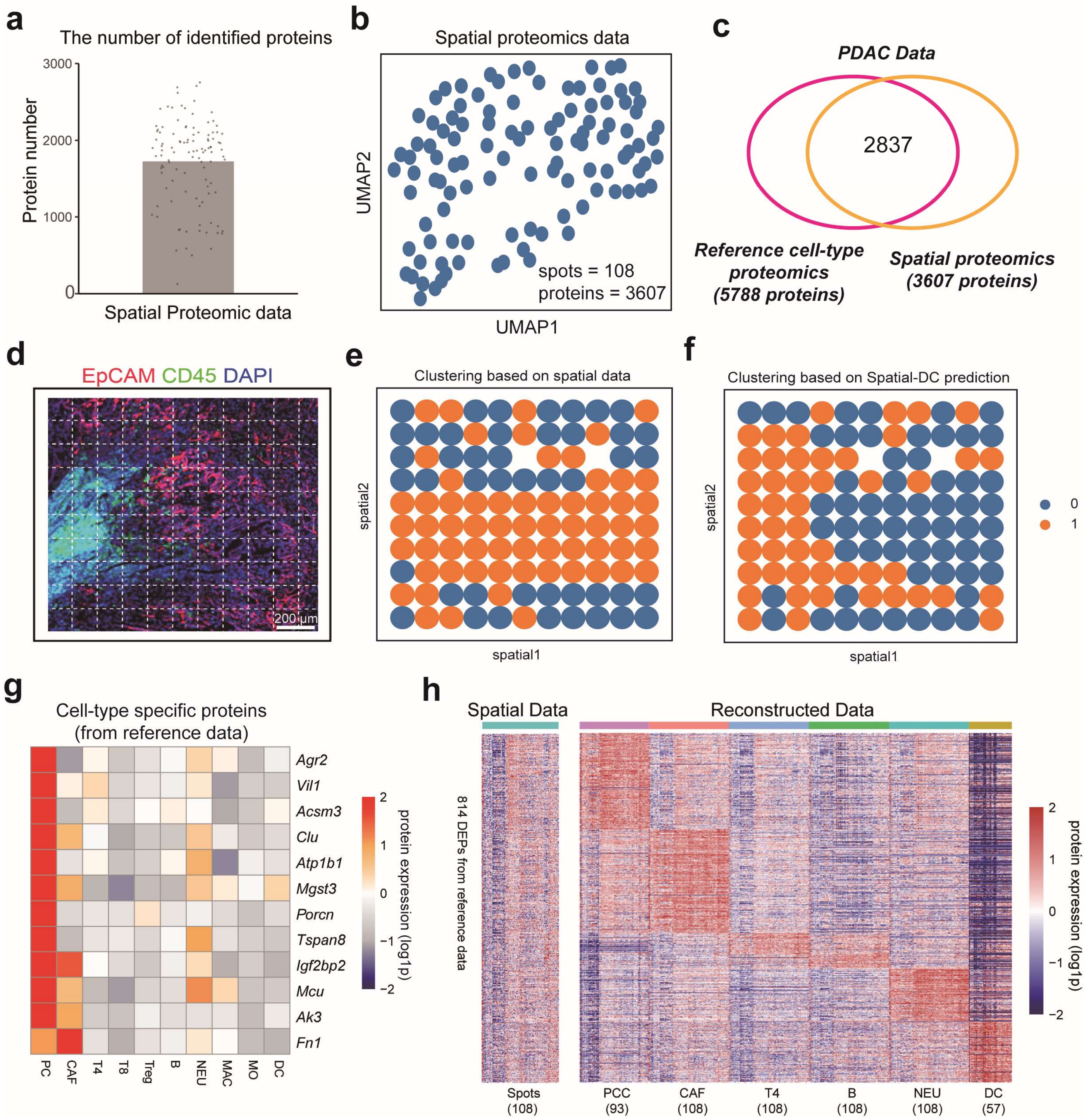


**Fig. S10 | Supplementary information on the application of Spatial-DC mouse PDAC data. a,** The numbers of identified proteins of 108 spots from mouse PDAC spatial proteomics data. The grey bar depicts the average number of identified proteins, while the scatter points represent those for individual spots. **b,** UMAP visualization of mouse PDAC spatial proteomics data. **c,** Venn diagram showing the compilation of mouse PDAC data, which consists of 2,854 proteins per spot and per sample, derived from the intersection of proteins in MS-based spatial proteomics data and reference single-cell-type proteomics data. **d,** Merged mIHC image depicting three-plex antibodies: *DAPI* for cell nucleus, *EPCAM* for cancer cells, and *CD45* for immune cells, sourced from mouse PDAC data (**Methods**). **e-f**, Comparison of spatial domain recognition using spot-level clustering and cell-type composition based-clustering with k-means clustering method (k = 2) (**Methods**). **g,** Heat map of the cell-type-specific proteins in **Fig. 5e**, using mean protein abundance from reference data. **h,** Comparison of 814 differentially expressed proteins (DEPs) from original spatial proteomics data (left) and reconstruction by Spatial-DC (right).

### Supplementary Notes

**Note S1. Pre-processing for the opensource and self-collected data in this study.**

**Synthetic NSCLC data.** The synthetic NSCLC data, utilized to generate **Fig. 2**, **Fig. S2 – Fig. S7**, comprises three parts: NSCLC spatial tissue structure data, NSCLC projection data, and NSCLC reference data. For the NSCLC spatial tissue structure data, the origin spatial tissue structures were accessible at https://nanostring.com/resources/smi-ffpe-dataset-lung13-data, which includes a total of 8 different samples from 5 NSCLC tissues. Among these, the tissue sample Lung 5-3 was utilized in subsequent simulations. For the NSCLC projection data, downloaded from the supplementary table of the referenced paper[2] at https://pubs.acs.org/doi/suppl/10.1021/acs.analchem.9b03349/suppl_file/ac9b03349_si_005.xlsx, the third sheet titled "02 - Processed sc protein data" was used, which contains the processed and log2-transformed protein intensity consisting of 1,225 proteins for 61 cells. In this study, the log2 transformation was reversed, and the missing values were replaced with zero. For the NSCLC reference data, the supplementary source data for "Fig.4b" from the referenced paper[1] was downloaded. This data also contains processed and log2-transformed protein intensity of 1,437 proteins for 108 cells. In this study, the log2 transformation was reversed, and no further processing steps were conducted on this data.

**Human palatine tonsil data.** The palatine tonsil data, utilized to generate **Fig. 3**, **Fig. S8**, comprises two parts: human palatine tonsil reference data[3] and human palatine tonsil spatial proteomics data[4]. The reference data was sourced from Zenodo (https://zenodo.org/records/8373756), which included processed CITE-seq data in form of Seurat objects. It consists of 192 proteins for 42,929 cells, collected from 15 donors. To minimize the impact of patient-related or batch effect-related biases, we selected the donor "BCLL-15-T", which contained 17,371 cells and originated from batch "BCLLATLAS_46", as our reference data. Rare cells were excluded. Ultimately, 19 cell types from 17,367 cells were retained for our analysis. The spatial data was downloaded from Gene Expression Omnibus (GSE213264) and includes 282 proteins for 2,492 spots. After manual curation for protein names, 174 proteins were found to be commonly quantified in both data. In this study, no external processing or imputation steps were applied to these data.

**Mouse brain coronal data.** The mouse brain coronal data, utilized to create **Fig. 4**, **Fig. S9**, incorporates two parts: mouse brain reference data[5] and mouse brain coronal spatial proteomics data[6]. The reference data, downloaded from MaxQB (http://maxqb.biochem.mpg.de/mxdb/project/show/P009), consists of 12,012 proteins across four primary mouse cell types, with three replicates for each cell type. For this data, we retained 10,734 proteins which have at least two valid values in one cell type. The spatial proteomics data, sourced from supplementary table of the referenced paper[6], includes 5,019 proteins from 208 spots. For this data, the log2 transformation was reversed. Missing values in both data were imputed using a strategy involving one sample's normal distribution of proteomic abundance using Perseus (down-shifted mean by 1.8 standard deviation and scaled by 0.3). In this study, there are 4,351 commonly quantified proteins in both data and no further processing steps were applied.

**Mouse PDAC data.** The mouse PDAC data, utilized to create **Fig. 5 and 6**, **Fig. S10**, incorporates two parts: mouse PDAC reference data[5] and mouse PDAC spatial proteomics data[6]. The reference data, downloaded from Zenodo (https://doi.org/10.5281/zenodo.13978420), consists of 5,900 proteins across 14 mouse PDAC cell types. In this study, we selected 10 primary mouse PDAC cell types, with each cell type collected from 5 $KP^{f/f}C$ mice. After filtering to ensure at least two valid values per cell type, we retained 5,788 proteins from 50 cells for the subsequent analysis. The self-collected spatial proteomics data contains 3,607 proteins from 108 spots. Missing values in both data were imputed using a strategy consistent with that used for the mouse brain coronal data. Notably, there are 2,837 proteins commonly quantified in both data and no additional processing steps were applied to these data in this study.

### Supplementary References


1. Woo, J. et al. High-throughput and high-efficiency sample preparation for single-cell proteomics using a nested nanowell chip. *Nat Commun* **12**, 6246 (2021).
2. Dou, M. et al. High-Throughput Single Cell Proteomics Enabled by Multiplex Isobaric Labeling in a Nanodroplet Sample Preparation Platform. *Anal. Chem.* **91**, 13119–13127 (2019).
3. Massoni-Badosa, R. et al. An atlas of cells in the human tonsil. *Immunity* **57**, 379-399.e18 (2024).
4. Liu, Y. et al. High-plex protein and whole transcriptome co-mapping at cellular resolution with spatial CITE-seq. *Nat Biotechnol* **41**, 1405–1409 (2023).
5. Sharma, K. et al. Cell type– and brain region–resolved mouse brain proteome. *Nat Neurosci* **18**, 1819–1831 (2015).
6. Ma, M. et al. In-depth mapping of protein localizations in whole tissue by micro-scaffold assisted spatial proteomics (MASP). *Nat Commun* **13**, 7736 (2022).

### Supplementary Table Titles

Table S1. The model configuration of Spatial-DC.

Table S2. Overview of synthetic data.

Table S3. Overview of real-world data.

Table S4. Assessment of Spatial-DC on synthetic NSCLC data.

Table S5. Application of Spatial-DC on human palatine tonsil data from antibody-based proteomics technology.

Table S6. Application of Spatial-DC on mouse brain coronal data from MS-based proteomics technology.

Table S7. Application of Spatial-DC for cell-type distributions on self-collected mouse PDAC data.

Table S8. Application of Spatial-DC for proteomic profiles and cell-cell communication on self-collected mouse PDAC data.

Table S9. Computational time (seconds) for Spatial-DC and eight state-of-the-art deconvolution methods.

Table S10. Overview of the known markers mentioned in this study.
